# Effects of clinically relevant radionuclides on the activation of a type I interferon response by radiopharmaceuticals in syngeneic murine tumor models

**DOI:** 10.1101/2024.07.10.602990

**Authors:** Caroline P. Kerr, Julia Sheehan-Klenk, Joseph J. Grudzinski, David P. Adam, Thanh Phuong T. Nguyen, Carolina A. Ferreira, Amber M. Bates, Won Jong Jin, Ohyun Kwon, Aeli P. Olson, Wilson Lin, Meredith Hyun, Justin C. Jagodinsky, Maria Powers, Raghava N. Sriramaneni, Paul A. Clark, Amanda G. Shea, Hansel Comas Rojas, Cynthia Choi, Christopher F. Massey, Luke M. Zangl, Anatoly N. Pinchuk, Eduardo Aluicio-Sarduy, KyungMann Kim, Jonathan W. Engle, Reinier Hernandez, Bryan P. Bednarz, Jamey P. Weichert, Zachary S. Morris

## Abstract

Radiopharmaceutical therapies (RPT) activate a type I interferon (IFN1) response in tumor cells. We hypothesized that the timing and amplitude of this response varies by isotope. We compared equal doses delivered by ^90^Y, ^177^Lu, and ^225^Ac *in vitro* as unbound radionuclides and *in vivo* when chelated to NM600, a tumor-selective alkylphosphocholine. Response in murine MOC2 head and neck carcinoma and B78 melanoma was evaluated by qPCR and flow cytometry. Therapeutic response to ^225^Ac-NM600+anti-CTLA4+anti-PD-L1 immune checkpoint inhibition (ICI) was evaluated in wild-type and stimulator of interferon genes knockout (STING KO) B78. The timing and magnitude of IFN1 response correlated with radionuclide half-life and linear energy transfer. CD8^+^/Treg ratios increased in tumors 7 days after ^90^Y- and ^177^Lu-NM600 and day 21 after ^225^Ac-NM600. ^225^Ac-NM600+ICI improved survival in mice with WT but not with STING KO tumors, relative to monotherapies. Immunomodulatory effects of RPT vary with radioisotope and promote STING-dependent enhanced response to ICIs in murine models.

**Teaser:** This study describes the time course and nature of tumor immunomodulation by radiopharmaceuticals with differing physical properties.

## Introduction

External beam radiation therapy (EBRT) has been shown to induce type I interferon (IFN1) responses by way of the cyclic GMP-AMP (cGAMP) synthase (cGAS) and downstream stimulator of interferon genes (STING) pathway (*1, 2*). IFN1 responses produce the inflammatory cytokine, IFNβ, which can then stimulate mature dendritic cells to cross-present tumor antigens and activate CD8^+^ T cells. In murine tumor models, EBRT-stimulated IFN1 responses can turn immunologically cold tumors into sites of activated T-cell infiltration and induce the further production of interferons by tumor cells (*2–4*). In preclinical models, EBRT to a single tumor site enhances responses to immune checkpoint inhibitors (ICIs) (*2, 5, 6*). This tumor regression is mediated by a cGAS/STING-dependent type I interferon (IFN1) response in cancer cells following sublethal doses of RT (8-12 Gy) (*7–9*). In certain preclinical studies, the murine tumor response to immunotherapy is abrogated by the presence of an unirradiated secondary tumor and restored upon irradiation of all tumor sites (*10, 11*).

In cases of radiographically occult or microscopic disease, radiation to all tumor sites using EBRT would require whole body treatment, resulting in lymphopenia and potentially prohibitive toxicities. Radiopharmaceutical therapy (RPT) offers systemic delivery of a therapeutic radionuclide to tumors using a small molecule, peptide, or antibody-based tumor-targeting vector. In murine and canine studies, beta (β) particle-emitting RPT has delivered immunomodulatory doses of radiation to tumors (2-5 Gy) without causing bone marrow suppression or systemic lymphopenia (*12–14*). A growing body of preclinical work has investigated the immunomodulatory effects of RPT on the tumor microenvironment (TME) and RPT agents in combination with immunotherapy (*15*).

In preclinical studies, tumor sites have been shown to selectively uptake and retain NM600, an alkylphosphocholine-DOTA chelator, that can be complexed with an array of radionuclides (^90^Y, ^177^Lu, ^225^Ac, ^64^Cu, ^86^Y) (*13*). By selective uptake of the radionuclide in every tumor location in the body, RPT has the potential to deliver radiation to all tumor sites in metastatic settings and, compared to EBRT approaches, would likely reduce toxicity and minimize systemic immunosuppression. We have confirmed the capacity of low-dose (∼2 Gy) ^90^Y-NM600 to activate a IFN1 response in B78 melanoma tumors in syngeneic mice (*16*). In a separate immunologically cold syngeneic tumor model, MOC2 head and neck squamous cell carcinoma (HNSCC), 2.5 or 12 Gy ^90^Y-NM600 led to IFN1 activation and induction of immune susceptibility markers (*Fas, Mhc1, Pdl1*, death receptor 5 (*Dr5*)) (*1*). Additionally, STING pathway activation was shown to be required for the therapeutic interaction between 1.85 MBq ^90^Y-NM600 and anti-CTLA4 in B16 melanoma, underscoring the critical role of this pathway (*16*).

Radionuclides that emit alpha (α; e.g., ^225^Ac, ^211^At, ^212^Pb, ^223^Ra) or β particles (e.g., ^90^Y, ^177^Lu), have different physical properties (e.g., emission type, linear energy transfer (LET), tissue range, half-life), which are hypothesized to confer different immunomodulatory effects. Few preclinical studies have compared α- and β-emitting radionuclide therapies directly. The high LET (50-230 keV/µm) of α-emitters results in a high density of ionization events, causing predominantly complex double-stranded DNA breaks (*17*). This short-range dose deposition may be effective in delivering dose to isolated or circulating tumor cells in addition to larger tumors. Conversely, the low LET (0-2 keV/µm) and longer-range emission (0.05-12 mm) of β-emitters results in a lower density of ionizing events as well as damaging but more readily repairable DNA breaks (*17*). Prior comparative radionuclide studies have noted different treatment outcomes of α-vs. β-emitting radionuclides conjugated to the same targeting moiety in murine tumors, some as monotherapies and others in combination with immune checkpoint blockade (*18–21*).

Using two poorly immunogenic murine tumor models, B78 melanoma and MOC2 HNSCC, we analyze and compare the effects of three distinct radionuclides, ^90^Y, ^177^Lu, and ^225^Ac (physical properties in **Table 1**), on the timing and magnitude of IFN1 response activation and immune susceptibility marker expression *in vitro* and *in vivo*. We hypothesized that ^225^Ac-NM600 (α-emitting, high LET, prolonged half-life, short range) may elicit an IFN1 response of greater magnitude and longer duration than ^90^Y- or ^177^Lu-NM600 (β-emitting, low LET, shorter half-life, long range). We further expected that ^225^Ac-NM600 and ICI would have a STING-dependent synergistic therapeutic effect.

**Table 1.** Physical properties of ^90^Y, ^177^Lu, ^225^Ac

| Isotope | Primary decay | Half-life (days) | Max range in tissue (mm) | Max Beta energy (keV) | Max Alpha energy (MeV) |
| --- | --- | --- | --- | --- | --- |
| <sup>90</sup>Y | Beta | 2.7 | 1.8 | 497 | NA |
| <sup>177</sup>Lu | Beta | 6.7 | 11.3 | 2284 | NA |
| <sup>225</sup>Ac | Alpha | 10 | <0.1 | NA | 5.83 |

## Results

### Magnitude of *in vitro* IFN1 response gene expression is greatest following ^225^Ac compared to ^90^Y and ^177^Lu

To demonstrate the effects of ^90^Y, ^177^Lu, or ^225^Ac on the magnitude and duration of a IFN1 response in tumor cells, we established *in vitro* dosimetry for each radionuclide (**Figure 1A-B**). *In vitro* RPT studies were used to differentiate tumor cell-specific immunomodulatory effects from *in vivo* immune effects of the gross tumor (including tumor cells, stroma, and infiltrating immune cells). In this experiment, four TLDs in the center of wells of a 6-well cell culture plate were exposed to a serial dilution of low-to moderate-dose free ^90^Y activity concentrations. The measured dose to the TLDs was within 15% error of the predicted dose to the TLDs estimated by GEANT4 Monte Carlo (**Figure 1B**). Additionally, Gafchromic film placed underneath the irradiated plate revealed a homogenous distribution of dose across the area of each well (**Figure 1B**). With this experimentally validated predictive dosimetry method, *in vitro* dosimetry was also determined for ^177^Lu and ^225^Ac.

**Fig. 1.**
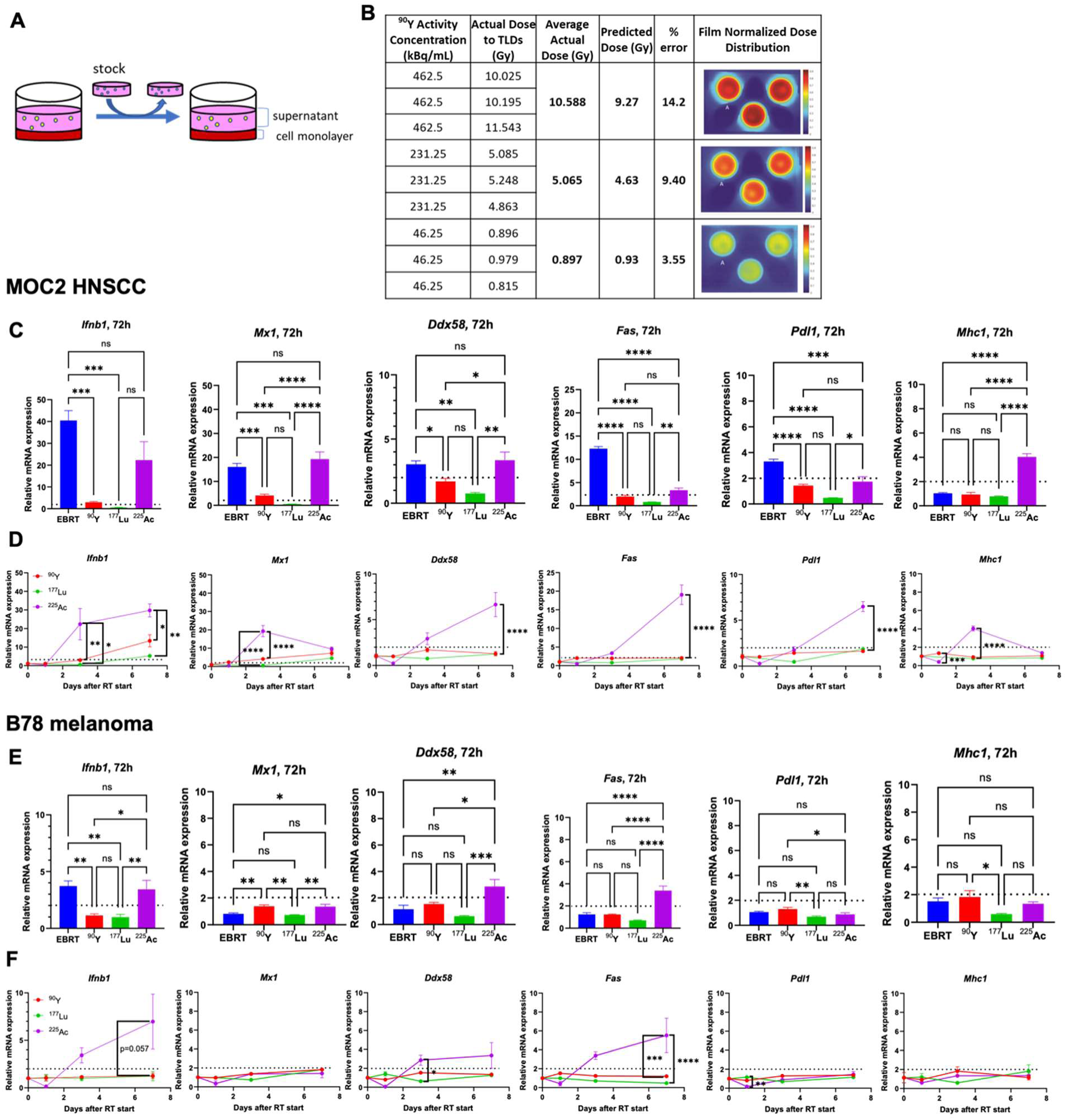
Magnitude of *in vitro* IFN1 response gene expression is greatest following ^225^Ac compared to ^90^Y and ^177^Lu. **(A)** Schematic of *in vitro* radionuclide therapy experiments showing unchelated radionuclide distributed in cell culture media overlaying adherent cell monolayer. **(B)** Four thermoluminescent dosimeters in center of well in 6-well plate exposed to 3 mL of 462.5, 231.25, or 46.25 kBq ^90^Y/mL RPMI, harvested after 10 days, and analyzed by the University of Wisconsin-Madison Radiation Calibration Laboratory (Calibration Cert #1664.01). These actual measurements were compared to predicted absorbed dose by GEANT4 Monte Carlo. Film normalized dose distribution to Gafchromic film under 6-well plate is shown. N=3 per ^90^Y activity concentration. **(C-F)** Cells were radiated with either 12 Gy (MOC2) or 4 Gy (B78) of EBRT, ^90^Y, ^177^Lu, or ^225^Ac and harvested 1, 3, or 7 days following RT. qPCR was used to quantify gene expression and is reported as fold changed normalized to untreated controls. Dotted line indicates a fold change of 2. N=3-6 per treatment group per timepoint. One-way **(C, E)** or two-way **(D, F)** ANOVA with Tukey’s HSD post hoc test was used to compare fold changes in expression between groups.

For *in vitro* studies, ^90^Y, ^177^Lu or ^225^Ac were added to culture media in activities estimated by the GEANT4 Monte Carlo method to deliver 12 Gy (MOC2) or 4 Gy (B78) to the cell monolayer. Expression of IFN1 response-associated genes (*Ifnβ1*, *Mx1*) was measured following radiation by each radionuclide. Additionally, we measured expression of *Ddx58*, the gene encoding RIG-I, a cytoplasmic double-stranded RNA sensor involved in both STING-dependent and STING-independent IFN1 production and downstream adaptive immune signaling following radiotherapy (reviewed by Zevini et al (*22*)). RIG-I-dependent IFN1 responses in tumor cells and dendritic cells have been demonstrated to be essential in the anti-tumor immune response following radiotherapy (*23, 24*). We also evaluated other immune susceptibility markers (*Fas, Pdl1, Mhc1*) previously evaluated in the context of EBRT and not directly related to STING and RIG-I signaling. Cell surface expression of these markers has been associated with susceptibility to immune-mediated elimination of tumor cells (*23, 25–31*). We delineated differences in timing and magnitude of mRNA expression of these genes in the setting of continuous radiation by an α- or β-emitting radionuclide (**Figure 1C-1F**). Compared to non-irradiated controls, ^225^Ac dosed to 12 Gy at infinity (7.4 kBq/mL) increased expression of each gene evaluated in MOC2 HNSCC cells (**Figure 1C**). The magnitude of gene expression following ^225^Ac was lower than that induced by EBRT at 72h following the start of irradiation; however, the difference was only significant for *Fas* and *Pdl1* (p < 0.0001, **Figure 1C**). For most genes (*Ifnb1, Ddx58, Fas, Pdl1*) evaluated, ^225^Ac maintained an increased expression level 7 days following the start of radiation exposure (**Figure 1D**).

For the low LET β-emitting radionuclides (^90^Y, ^177^Lu), *Ifnb1* and *Mx1* expression were increased relative to control at 7 days post radiation therapy (RT) start; however, the magnitude of *Ifnb1* expression was significantly lower than that following ^225^Ac **(Figure 1D**). In MOC2 cells following 12 Gy doses of ^90^Y and ^177^Lu at infinity, *Ddx58*, *Fas*, *Pdl1*, and *Mhc1* expression were not upregulated (**Figure 1D**). IFN1 activation following ^90^Y, ^177^Lu, and ^225^Ac in B78 melanoma cells was evaluated at a lower dose (4 Gy), previously shown to be immunomodulatory for ^90^Y-NM600 + ICI responses in B78 melanoma (*16*). However, given that this lower dose is outside the optimal 8-12 Gy IFN1 activating range following EBRT (*9*), gene expression effects were tempered for all forms of irradiation, with minimal to no upregulation of gene expression following EBRT, ^90^Y, nor ^177^Lu (**Figure 1E-F**). In contrast to other radiation modalities, α-emitting ^225^Ac prescribed to 4 Gy at infinity increased expression of *Ifnb1*, *Ddx58*, and *Fas* until 7 days following RT start (**Figure 1F**).

Also, in MOC2 HNSCC cells, we evaluated *in vitro* expression of these IFN1 response-associated genes following treatment with the Auger-emitting radionuclides, ^58m^Co and ^119^Sb, which have differing half-lives (^58m^Co: 9h, ^119^Sb: 38h) (*32, 33*). For these radionuclides, a serial dilution of activities was added to the cell culture media, and mRNA was quantified at 24 hours, 72 hours, and 7 days. After 72 hours, expression of *Ifnb1* increased in the presence of 370 or 37 kBq, but not 3.7 kBq, of ^58m^Co (**Figure S2A**). Following 370 or 37 kBq ^119^Sb irradiation, upregulated *Ifnb1* and *Mx1* expression in MOC2 cells was observed at 72h and was similar in magnitude to that following ^58m^Co (**Figure S2B)**.

The activation of IFN1 responses across five radioisotopes with different particle emissions suggests that radiation in many forms leads to IFN1 response activation, and the magnitude and duration of this response may be influenced by the LET and dose rate of the emitted radiation.

### Timing and magnitude of IFN1 responses may correlate with radiation dose rate and LET

To clarify the relationship of the radiation dose rate and LET to IFN1 response, the effects of dose rate and absorbed dose of each radionuclide were evaluated in isolation by performing EBRT fractionation and dose equivalent studies, respectively (**Figure 2A-C**). EBRT fractionation (EBRT-fr; orange on graphs) studies involved irradiating cells with EBRT every 24 hours with the corresponding dose delivered by ^90^Y, ^177^Lu, or ^225^Ac when prescribed to a 12 Gy at infinity dose (**Figure 2A**, **Table S2**). EBRT-fr irradiation mimics dose rate of a radioisotope, while maintaining the critical difference of allowing a 24h period of DNA repair between fractions, as opposed to continuous irradiation by a radioisotope. EBRT dose equivalent (EBRT-eq; black on graphs) studies involved EBRT on day 0 only with the dose delivered by each radioisotope at each gene expression harvest timepoint (**Figure 2B-C**, **Table S2**). The EBRT-eq group addresses whether the differed IFN1 response activation observed following ^90^Y, ^177^Lu, ^225^Ac can be attributed to varied dose accumulation over the time course of radionuclide decay as opposed to the lower cumulative dose at each harvest timepoint alone, given the *in vitro* dose prescription to infinity.

**Fig. 2.**
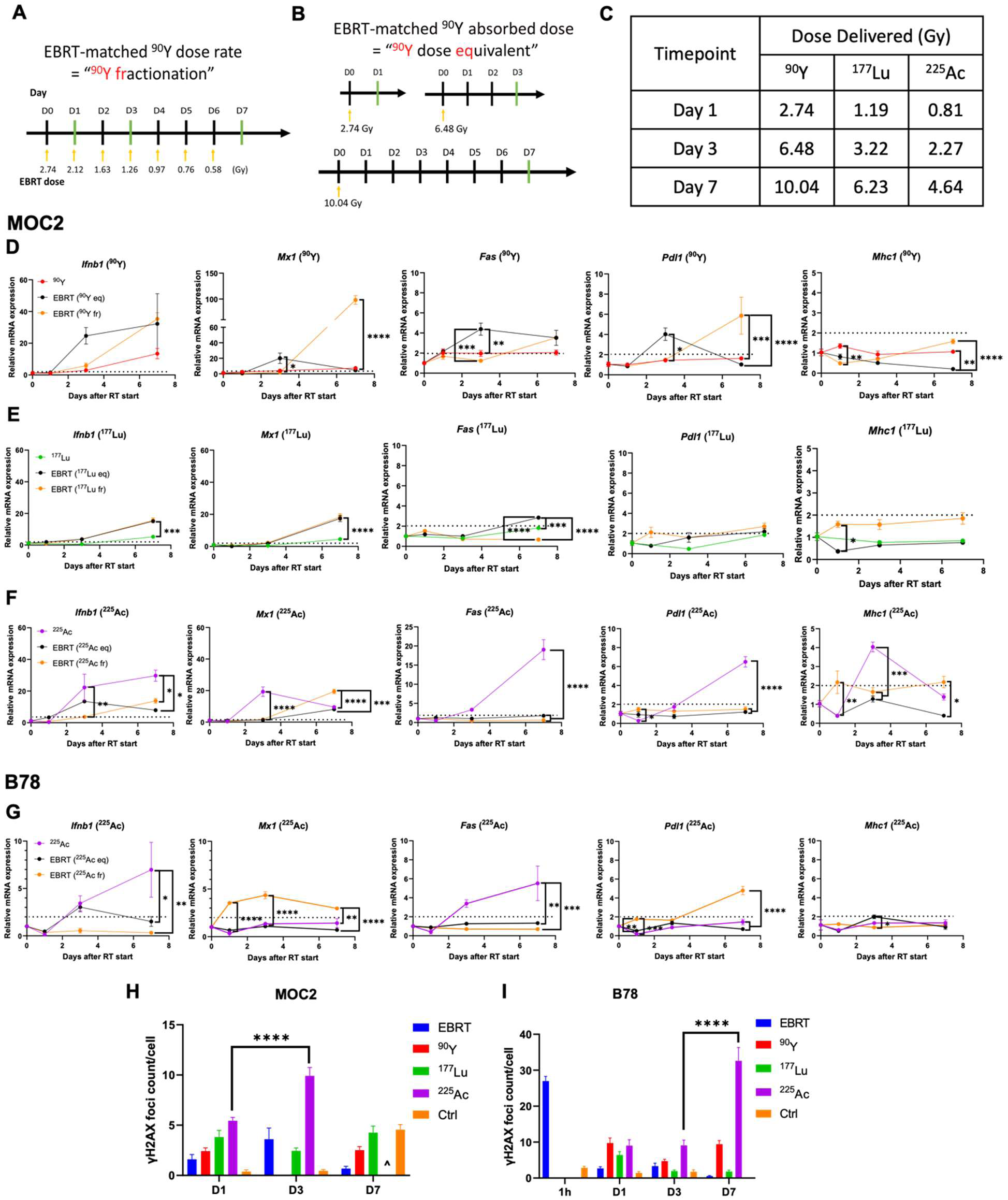
Timing and magnitude of radionuclide-induced type I IFN responses may correlate with radiation dose rate and LET. **(A-B)** Schematic of EBRT dosing regimens corresponding to 12 Gy ^90^Y dosed to infinity. Yellow arrows indicate EBRT dose administered, green ticks indicate cell harvest timepoint. **(C)** Table showing dose delivered by ^90^Y, ^177^Lu, or ^225^Ac at each cell harvest timepoint when prescribed to deliver 12 Gy at an infinite timepoint. **(D-I)** MOC2 (12 Gy) and B78 (4 Gy) cells treated with EBRT, ^90^Y, ^177^Lu, or ^225^Ac dose according to treatment regimens in Tables S1 and S2, and harvested or fixed 1, 3, or 7 days following RT. **(D-G)** qPCR was used to quantify gene expression and is reported as fold changed normalized to untreated controls. **(H-I)** Cells fixed with 4% paraformaldehyde, probed with immunofluorescent anti-γH2AX antibody, and analyzed with ImageJ. γH2AX foci were not quantified for the MOC2 EBRT 1h group due to the high prevalence of apoptotic rings as opposed to nuclear foci nor for the MOC2 ^225^Ac day 7 group due to the low number of nuclei present; missing value indicated by “^”. MOC2 qPCR: n=3-6, repeated; B78 qPCR: n=5-6, repeated. γH2AX immunofluorescence: n=50 cells quantified. Two-way ANOVA with Tukey’s HSD post hoc test was used to compare fold changes in expression, foci counts between groups.

Cells treated every 24h with EBRT-matched ^90^Y/^177^Lu/^225^Ac dose rates (EBRT-fr; orange) resulted in IFN1 response time courses more closely mimicking radionuclide-induced responses than the other EBRT regimens (**Figure 2D-G**; **S3**). The magnitude of the radionuclide-induced *Ifnb1* expression was lower than EBRT fractionation-induced expression on day 7 for low LET ^90^Y and ^177^Lu (red/green vs. orange, **Figure 2D-E**); but higher in magnitude than the dose rate-mimicking EBRT-fr condition for high LET ^225^Ac (purple vs. orange, **Figure 2F-G**). *Fas* and *Pdl1* expression showed a similar pattern to *Ifnb1*, where the magnitude of gene expression was higher for the ^225^Ac group than the corresponding EBRT-fr group, and lower for ^90^Y as compared to the EBRT-fr group (**Figure 2D**, **F**).

Because cytosolic double-stranded DNA leads to cGAS/STING activation following radiation damage (*7*), we measured repair of double-stranded DNA breaks with immunofluorescent staining of γH2AX foci. γH2AX foci counts/cell increased significantly and accumulated over time following treatment with ^225^Ac, but not with ^90^Y or ^177^Lu for both MOC2 HNSCC and B78 melanoma (**Figures 2H-I**; **S4**). However, ^225^Ac induced a comparable amount of double-stranded DNA damage in B78 cells as EBRT but over a longer time course (7 days vs. 1 h; **Figure 2I**). These findings correlated with the gene expression magnitude differences observed in IFN1 activation between EBRT-fr regimens and the unconjugated radionuclides (^90^Y, ^177^Lu < EBRT < ^225^Ac). Overall, ^225^Ac induced more double-stranded DNA damage over time than ^90^Y and ^177^Lu, and IFN1 response gene expression of higher magnitude but similar time course to EBRT absorbed dose-matched and dose-rate-matched counterparts, respectively. β-emitting ^90^Y and ^177^Lu therapy led to lower magnitude but similar time course IFN1-response gene expression than EBRT counterparts. Auger electron-emitting ^58m^Co and ^119^Sb therapy showed earlier peaks in *Ifnb1* upregulation than ^90^Y and ^177^Lu, but not high LET ^225^Ac (**Figures 1D**; **S2**).

### Imaging, uptake, and dosimetry of ^177^Lu-NM600 and ^225^Ac-NM600 in MOC2 HNSCC, B78 WT and STING KO melanoma

To compare immunomodulatory effects of ^90^Y-NM600, ^177^Lu-NM600 and ^225^Ac-NM600 *in vivo*, image-based radiation dosimetry studies were performed (NM600 chemical structure: **Figure 3A**). Image-based dosimetry for ^90^Y-NM600 in mice bearing B78 (*16*) and MOC2 (*1*) tumors using ^86^Y-NM600 as a theranostic pair was previously obtained and reported. For ^177^Lu-NM600, serial SPECT/CT images were obtained at various time points following ^177^Lu-NM600 administration; maximum intensity projections demonstrating selective tumor uptake are shown in **Figure 3B,D**. Toxicity assessments for ^90^Y-NM600 and ^177^Lu-NM600 have been previously reported (*34, 35*). Absorbed dose estimates in Gy/MBq for ^177^Lu-NM600 in MOC2 or B78 tumor-bearing mice are shown in **Figure 3C**,**E**. Longitudinal SPECT/CT images were used to estimate dose to tumor and normal tissues using our Monte Carlo voxel-based dosimetry method (*36*).

**Fig. 3.**
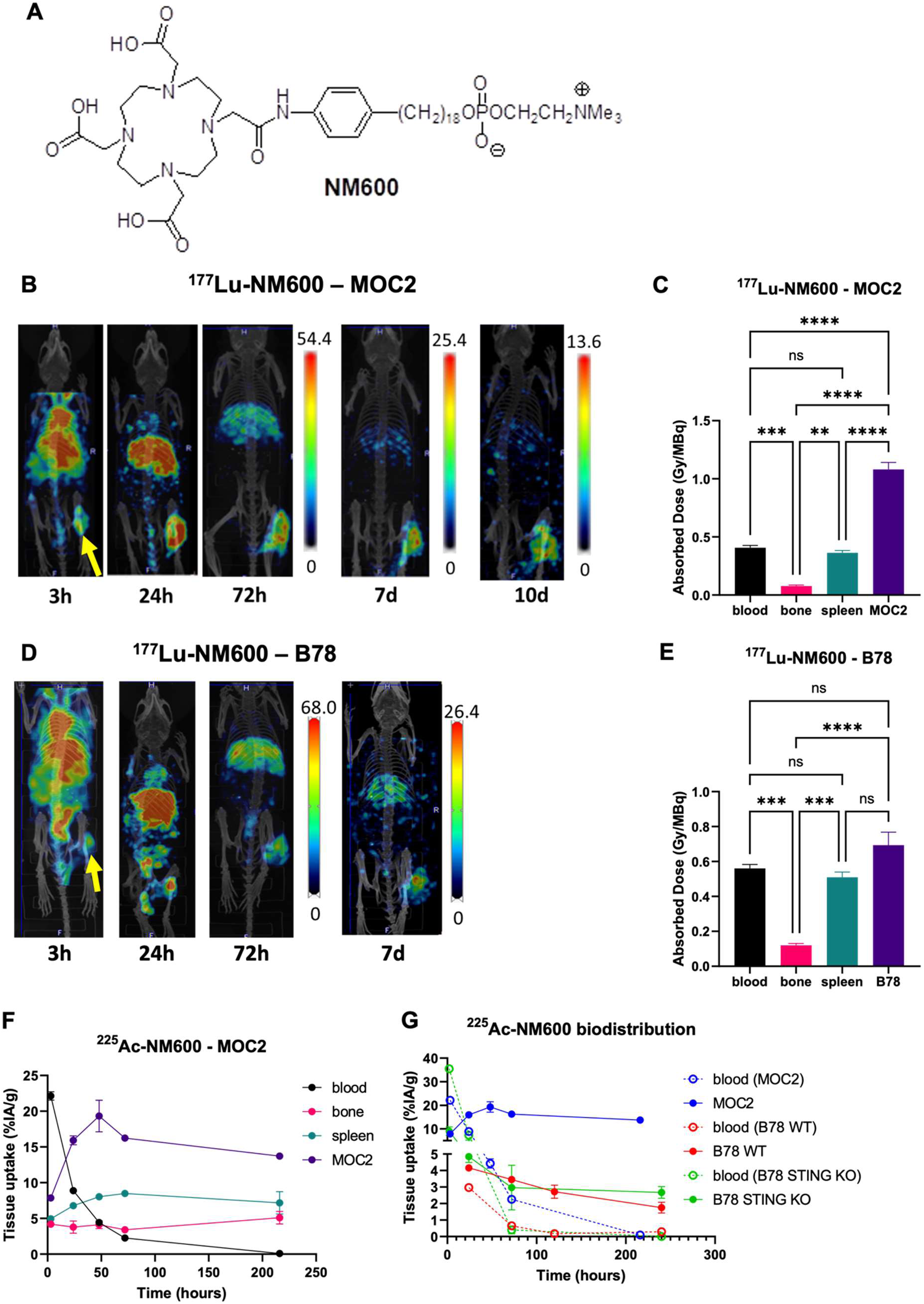
Imaging, uptake, and dosimetry of ^177^Lu-NM600 and ^225^Ac-NM600 in MOC2 HNSCC, B78 WT and STING KO melanoma. **(A)** Chemical structure of NM600. **(B)** Maximum intensity projections (MIPs) of serial SPECT/CT images 3h, 24h, 72h, 168h, and 240h post-injection of 18.5 MBq ^177^Lu-NM600 for a MOC2 HNSCC tumor-bearing mouse (tumor indicated by arrow). **(C)** Tissue dosimetry was performed using Monte Carlo methods (n=3). **(D)** Maximum intensity projections (MIPs) of serial SPECT/CT images 3h, 24h, 72h, and 168h post-injection of 18.5 MBq ^177^Lu-NM600 for a B78 melanoma tumor-bearing mouse (tumor indicated by arrow). **(E)** Tissue dosimetry was performed using Monte Carlo methods (n=3). **(F)** *Ex vivo* biodistribution of ^225^Ac-NM600 in MOC2-tumor bearing mice. **(G)** *Ex vivo* tissue uptake in blood and MOC2, B78 WT, or B78 STING KO flank tumor following tail vein injection of 9.25 kBq ^225^Ac-NM600. N=3/timepoint; MOC2 timepoints: 3h, 24h, 48h, 72h, 216h; B78 WT timepoints: 24h, 72h, 120h, 240h; B78 STING KO timepoints: 2h, 24h, 72h, 240h. **(B, D)** One-way ANOVA with Tukey’s HSD post hoc test used to compare absorbed doses (Gy/MBq) between tissues.

MOC2 and B78 tumors received the highest dose of all tissues, at mean normalized doses of 1.08 Gy/MBq and 0.81 Gy/MBq, respectively. Absorbed dose for the MOC2 tumor (mean +/-standard deviation = 1.08 +/-0.11 Gy/MBq) was significantly higher than immune organs (blood: 0.41 +/-0.03, bone: 0.08 +/- 0.02, spleen: 0.36 +/- 0.04 Gy/MBq), corroborating selective tumor uptake of ^177^Lu-NM600 (**Figure 3C**). B78 melanoma (0.81 +/- 0.07 Gy/MBq) showed lower tumor uptake than MOC2, reflective of tumor model differences observed with the ^86^Y-NM600 analog as well. ^177^Lu-NM600 was selective for B78 however, demonstrating higher uptake in the tumor than immune organs and significantly higher tumor absorbed dose than bone absorbed dose (blood: 0.56 +/0 0.04, bone: 0.12 +/- 0.02, spleen: 0.51 +/- 0.05 Gy/MBq).

Serial *ex vivo* biodistribution data were used to estimate dose to the tumor and normal tissues for ^225^Ac-NM600. ^225^Ac-NM600 tissue uptake over time for mice bearing either MOC2, B78 WT, or B78 STING KO tumors is shown in **Figures 3C-D**. Tumor absorbed doses of 1.31, 0.10, and 0.12 Gy/kBq were estimated for MOC2, B78 WT, and B78 STING KO tumors, respectively. ^225^Ac-NM600 was cleared from the blood within 72 hours of injection and was retained in the tumor at all time points measured (up to 9 days post-injection; **Figure 3F-G**). All doses of RPT investigated *in vivo* were determined by radionuclide- and tumor-model-specific dosimetry and were well tolerated.

### *In vivo* type I IFN response activation is prolonged following ^90^Y-, ^177^Lu-, ^225^Ac-NM600 compared to tumor-targeted or whole-mouse EBRT

We next investigated the effects of radioisotope physical properties on the *in vivo* IFN1 response. In mice bearing MOC2 flank tumors, we irradiated either the tumor only using lead shielding (tumor EBRT), the whole mouse (whole mouse radiation therapy; WMRT), or systemically with ^90^Y-, ^177^Lu-, or ^225^Ac-NM600, administered on day 1 at a tumor dose prescription of 12 Gy at infinity (**Figure 4A**). WMRT was included as a control for a clinically viable alternative approach to delivering low dose radiation to all tumor sites using EBRT in settings of potential radiographically occult metastatic disease. On days 3, 7, 14, and 21, tumors were harvested and disaggregated for gene expression analyses (**Figure 4B**). We observed similar trends to our *in vitro* experiments, in which tumor EBRT and WMRT led to earlier, higher magnitude upregulation of *Ifnb1*, *Mx1*, *Pdl1*, and *Mhc1* than RPT (**Figure 4C**). Longitudinal gene expression results following 12 Gy tumor-targeted EBRT are shown in **Figure S5**. Consistent with radiation dose being deposited slowly and continuously over a period of radionuclide decay, *Ifnb1* expression was either maintained at an elevated level compared to control tumors between day 3 and day 7 (^90^Y-, ^225^Ac-NM600) or increased over the same period (^177^Lu-NM600) for the RPT groups (**Figure 4C**). This sustained IFN1 activation may result in a favorable TME for immunotherapy response with prolonged dendritic cell recruitment (*37–39*) in RPT-treated tumors as compared to EBRT. For ^90^Y-, ^177^Lu-, and ^225^Ac-NM600 in MOC2 tumors, *Ifnb1* remained increased through day 7 for ^90^Y- and ^225^Ac-NM600 and through day 14 for ^177^Lu-NM600 (**Figure 4D**). *Ifnb1* and IFN1-stimulated gene *Mx1* returned to baseline levels by day 21 following RPT (**Figure 4D**).

**Fig. 4.**
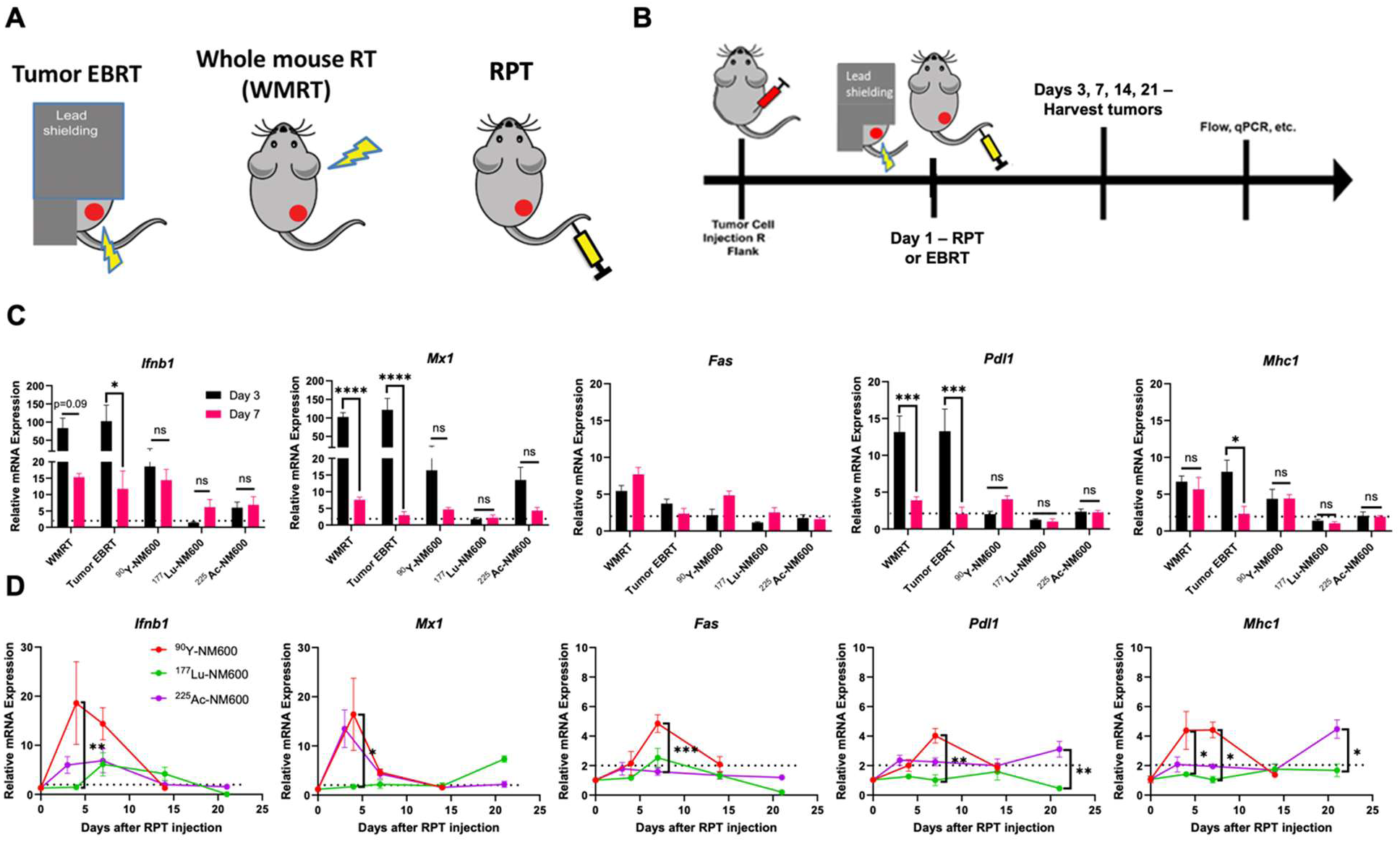
*In vivo* type I IFN response activation is prolonged following ^90^Y-, ^177^Lu-, ^225^Ac-NM600 compared to tumor-targeted or whole mouse external beam radiation therapy (EBRT). **(A)** Schematic of radiation treatments administered for *in vivo* experiments. **(B)** Experimental design for *in vivo* qPCR and flow cytometry experiments. MOC2 tumors were grown to ∼200 mm^3^ and radiated with either 12 Gy tumor targeted EBRT, whole mouse radiation therapy (WMRT), or ^90^Y-, ^177^Lu- or ^225^Ac-NM600 prescribed to deliver a tumor dose of 12 Gy at infinity. Tumors were harvested 3, 7, 14, or 21 days following RT administration. **(C-D)** qPCR was used to quantify gene expression and is reported as fold changed normalized to untreated controls. N=3-6 mice per treatment group per timepoint. **(C)** Unpaired two-tailed t-test used to compare fold changes between day 3 and day 7 for each group. **(D)** Two-way ANOVA with Tukey’s HSD post hoc test was used to compare fold changes in expression between groups.

### 90Y-, ^177^Lu, ^225^Ac-NM600 modulate tumor immune cell composition in the TME

Type I interferons following EBRT are critical for enhancing dendritic cell cross-priming of CD8^+^ T cells (*40*) and recruiting and increasing effector molecule production of CD8^+^ T cells in the TME (*37*). We evaluated these downstream effects of IFN1 signaling on tumor infiltrating lymphocyte (TIL) populations following RPT. Using the same MOC2-bearing mice as for the qPCR experiments above, we investigated the immune cell composition of host immune organs (spleen, bone marrow, blood, tumor-draining lymph node) and disaggregated tumor tissue compared to untreated control mice with matched size tumors by flow cytometry. In these tissues, CD45^+^, CD8^+^, regulatory T cell (Treg), CD8^+^/Treg ratio, and %PD1^+^ CD8^+^ T cells are reported at days 3, 7, 14, and 21 following RT on day 1 in **Figure 5**. In **Figure S6**, CD4^+^ T cells, NK cells, CD11b^+^ cells, M1- and M2-like macrophages, and dendritic cells are reported for the same treatment groups, tissues, and timepoints. The CD8^+^/Treg ratio significantly increased (p < 0.01) relative to control mice in the tumor only on day 7 after ^90^Y- and ^177^Lu-NM600 and on day 21 after ^225^Ac-NM600 as compared to spleen, bone marrow, blood, and tumor-draining lymph node (**Figure 5C-E**). Notably, a significant change in CD8^+^ frequency compared to control was not observed in the tumor for any RPT, but tumor Treg percentage was decreased relative to control (ratio < 0.5) for ^90^Y- and ^177^Lu-NM600 on day 7 and ^225^Ac-NM600 on day 21 (**Figure 5C-E**). In the ^90^Y-NM600 and tumor-targeted EBRT groups only, the percentage of tumoral CD45^+^ cells out of live cells relative to control mice increased significantly over other immune organs (**Figure 5B-C**; ^90^Y-NM600: all other immune organs, p < 0.001; tumor EBRT: blood, p < 0.05). This finding suggests that higher-dose-rate radiation may influence immune cell trafficking and infiltration more than lower-dose-rate radiation, consistent with a higher magnitude IFN1 response (**Figure 4C-D**). In contrast, lower dose rate radiation may modulate the composition of immune cells in the TME without changing total number of live immune cells. Each RPT, but not tumor EBRT, significantly increased the frequency of CD11b^+^ cells in the tumor draining lymph node relative to the tumor and other immune organs (**Figure S6C-E**). This finding warrants further investigation regarding subtype and function of these lymph node-residing myeloid cells. Overall, these results demonstrated unique impacts of ^90^Y-, ^177^Lu-, and ^225^Ac-NM600 on the TME.

**Fig. 5.**
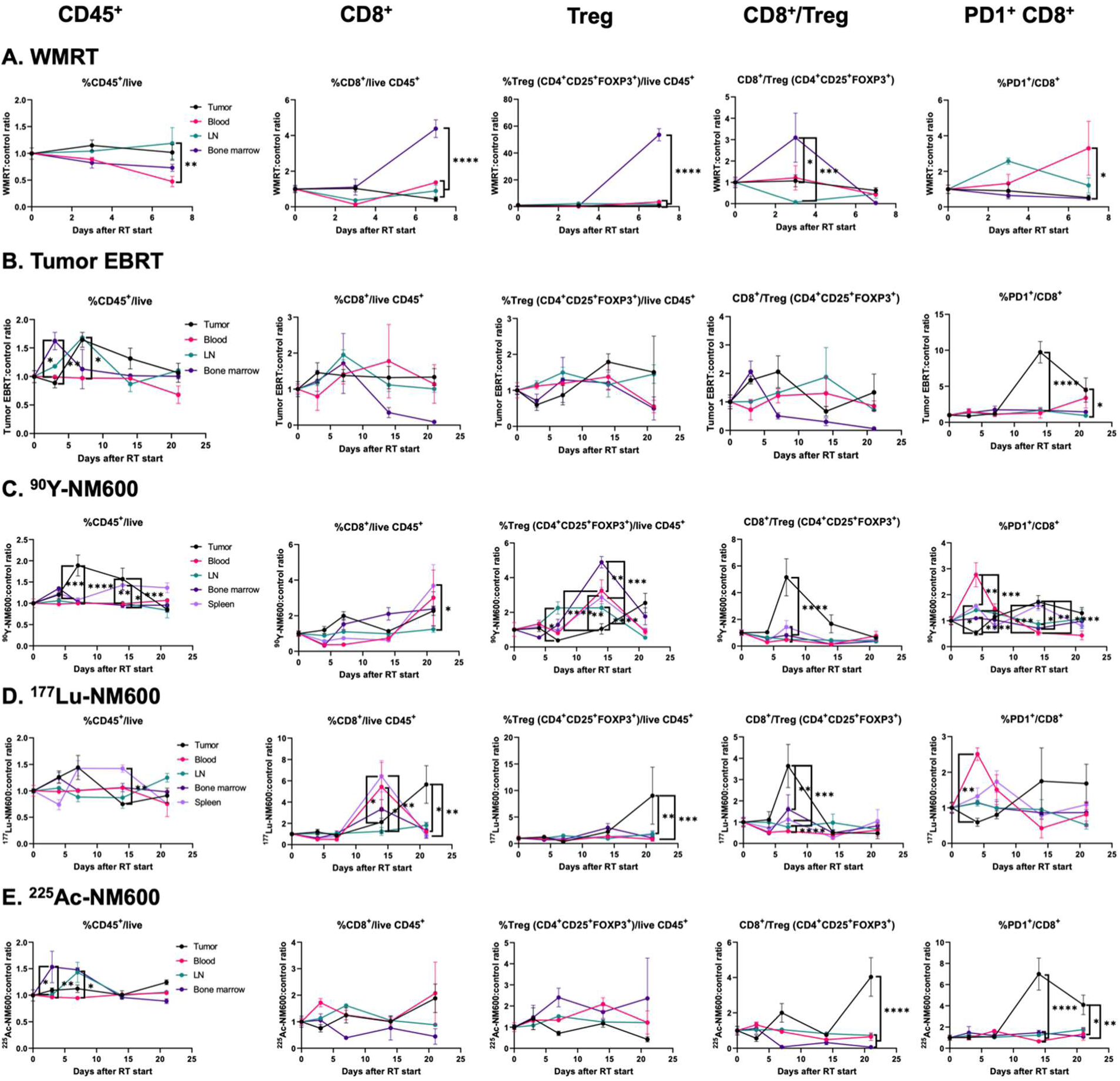
^90^Y-, ^177^Lu, ^225^Ac-NM600 modulate tumor immune cell composition in the tumor microenvironment. **(A-E)** From same mice as Figure 4, flow cytometry analyses of tumor, blood, tumor-draining lymph node, bone marrow, and spleen immune cell infiltrates [total immune cells (CD45^+^), T effector cells (CD8^+^), regulatory T cells (Tregs; CD4^+^CD25^+^FOXP3^+^), CD8^+^/Treg ratio, and PD1^+^ CD8^+^ T cells] as a percent of total live cells normalized to mean of control (no treatment) is shown at 3, 7, 14, and 21 days after RT administration in MOC2 HNSCC. N=4-6 per treatment group per timepoint. Two-way ANOVA with Tukey’s HSD post hoc test was used to compare ratios between tissue types.

### Cooperative therapeutic interaction between ^225^Ac-NM600 + ICI is STING dependent

STING is responsible for the synergistic response to EBRT + ICI (*9*) and ^90^Y-NM600 + ICI (*16*), but STING’s requirement for ^177^Lu- or ^225^Ac-based RPT + ICI combination therapies is unknown. Following 4 Gy ^225^Ac-NM600 or EBRT, *Ifnb1* expression was upregulated *in vitro* in B78 WT and STING KO cell lines (**Figure 6B-C**, **S7**), which suggested involvement of a STING-independent mechanism of IFN1 activation. Following 4 Gy ^225^Ac-NM600 but not EBRT, *Ddx58* (RIG-I) transcription was upregulated *in vitro* in both B78 WT and STING KO cell lines (**Figure 6D-E**, **S7**). While Patel et al. previously reported that effects of low dose ^90^Y-NM600 in enabling response to ICIs in poorly immunogenic tumors were STING dependent (*16*), this has not been established for other radioisotopes. Given our observation that ^225^Ac upregulates RIG-I, which can activate IFN1 by an alternative pathway to STING, we tested whether ^225^Ac-NM600 could promote ICI response and whether this effect would be STING-dependent.

**Fig. 6.**
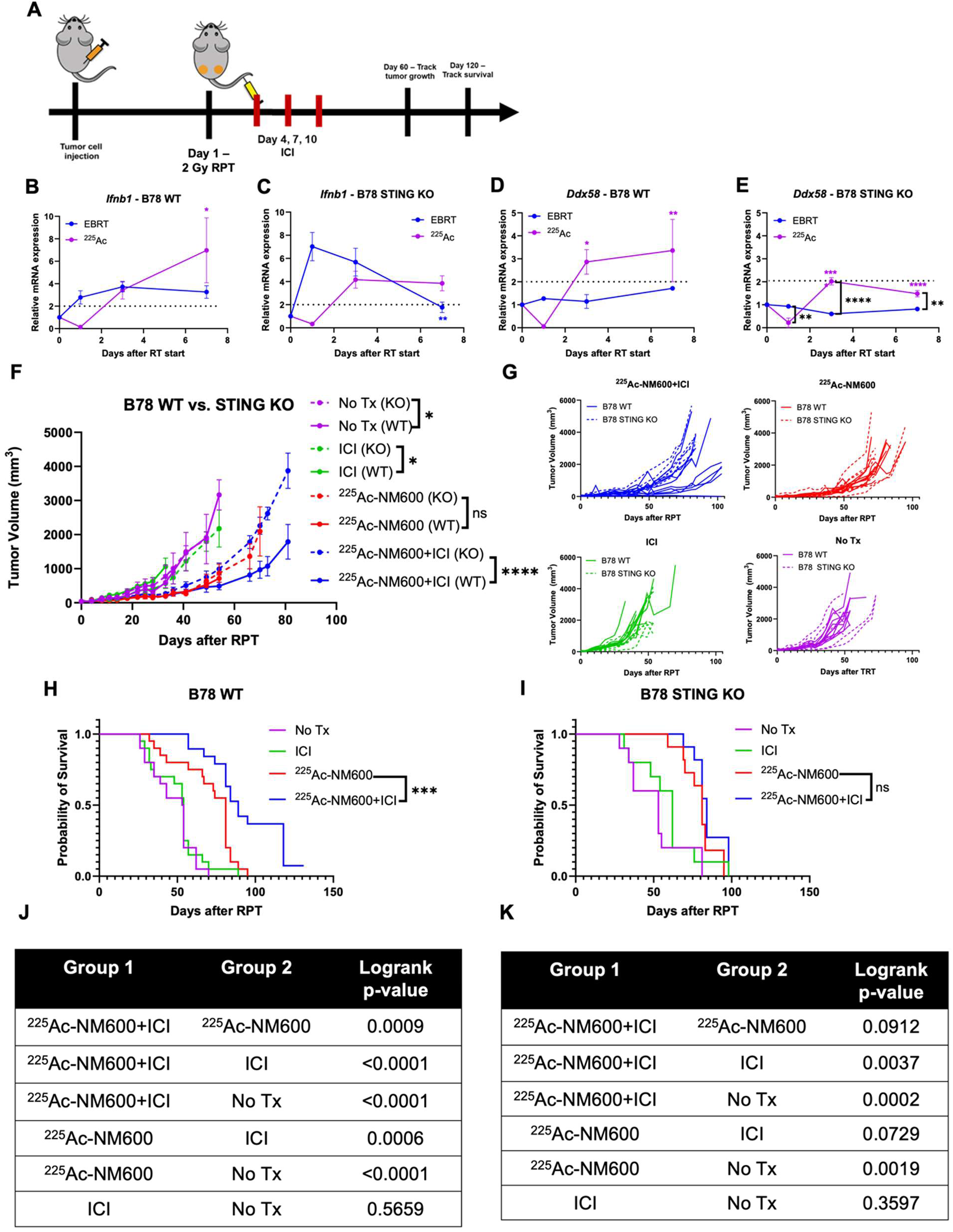
Cooperative therapeutic interaction between ^225^Ac-NM600 + ICI is STING dependent. **(A)** Treatment scheme for *in vivo* therapy studies. **(B-E)** *In vitro* qPCR following 4 Gy EBRT or ^225^Ac. qPCR was used to quantify gene expression and is reported as fold changed normalized to untreated controls. qPCR: n=4-6 per treatment group per timepoint. Two-way ANOVA with Tukey’s HSD post hoc test was used to compare fold change in expression between cell lines. Colored asterisks correspond to comparisons to day 1 for EBRT (blue) or ^225^Ac (purple). Black asterisks correspond to comparisons between treatment groups at the indicated timepoint. **(F-K)** B78 wild-type (WT) and STING knockout (STING KO) two-tumor bearing mice were randomized to untreated control (No Tx), dual anti-CTLA4 and anti-PD-L1 (ICI), 2 Gy ^225^Ac-NM600 (18.5 kBq), or combination ^225^Ac-NM600 and ICI. Combination ^225^Ac-NM600+ICI reduces tumor growth rate in B78 WT tumors compared to B78 STING KO tumors **(F-G)** and prolongs survival in B78 WT, but not B78 STING KO, tumors compared to ^225^Ac-NM600 monotherapy **(H-K)**. B78 WT tumor measurements (TMs) all treatment groups: n=10; B78 STING KO TMs: ^225^Ac-NM600+ICI: n=6, ^225^Ac-NM600: n=6, ICI: n=5, No Tx: n=5; B78 WT aggregate of two survival studies: ^225^Ac-NM600+ICI: n=19, ^225^Ac-NM600: n=20, ICI: n=20, No Tx: n=20; B78 STING KO survival studies: ^225^Ac-NM600+ICI: n=11, ^225^Ac-NM600: n=11, ICI: n=10, No Tx: n=10. Log-rank test was used to compare survival.

We tested the capability of ^225^Ac-NM600 to convert a poorly immunogenic tumor (B78 melanoma) to a tumor that responds to ICIs (**Figure 6A**). Combination ^225^Ac-NM600 + dual ICI (anti-CTLA4 + anti-PD-L1) significantly improved overall survival over ^225^Ac-NM600 (p = 0.0009) or dual ICI therapy (p < 0.0001) in mice bearing wild-type B78 melanoma flank tumors (**Figure 6H**, **J**). In mice bearing B78 STING KO tumors, there was not a statistically significant survival benefit from ^225^Ac-NM600 + dual ICI over ^225^Ac-NM600 monotherapy (**Figure 6I**, **K**; p = 0.0912). Despite *Ifnb1* production in the B78 STING KO cell line following ^225^Ac-NM600, STING expression in the tumor was necessary for the cooperative therapeutic interaction between ^225^Ac-NM600 and dual ICI (**Figure 6F-K**). The tumor growth curves show an equivalent response to the ^225^Ac-NM600 + ICI combination therapy early in the study (**Figure 6F-G**), but ultimately the STING KO cell line did not display the durability of tumor growth delay of the B78 melanoma WT cell line.

### 225Ac-NM600+ICI extends greater survival benefit than ^177^Lu-, ^90^Y-NM600+ICI in two-tumor B78 melanoma model

Given differences in type I interferon response gene upregulation and tumor infiltrate composition following ^225^Ac-NM600 as compared to ^90^Y- and ^177^Lu-NM600, the comparative therapeutic efficacy of each RPT in combination with dual ICI was evaluated in a two-tumor model (**Figure 7**, **Tables S7-8**). The two tumor model is more difficult to cure than a single flank tumor given the greater initial disease burden, but is a more representative model for the metastatic disease setting that is the intended clinical application of RPT+ICI. Combination ^225^Ac-NM600+ICI reduces tumor growth rate in B78 WT tumors compared to ^90^Y-NM600+ICI, ^177^Lu-NM600, EBRT+ICI, and No Tx groups (**Figure 7B-C**, **Table S7**). The longest median survival was measured following combination ^225^Ac-NM600+ICI (median survival: 81 days).

**Fig. 7.**
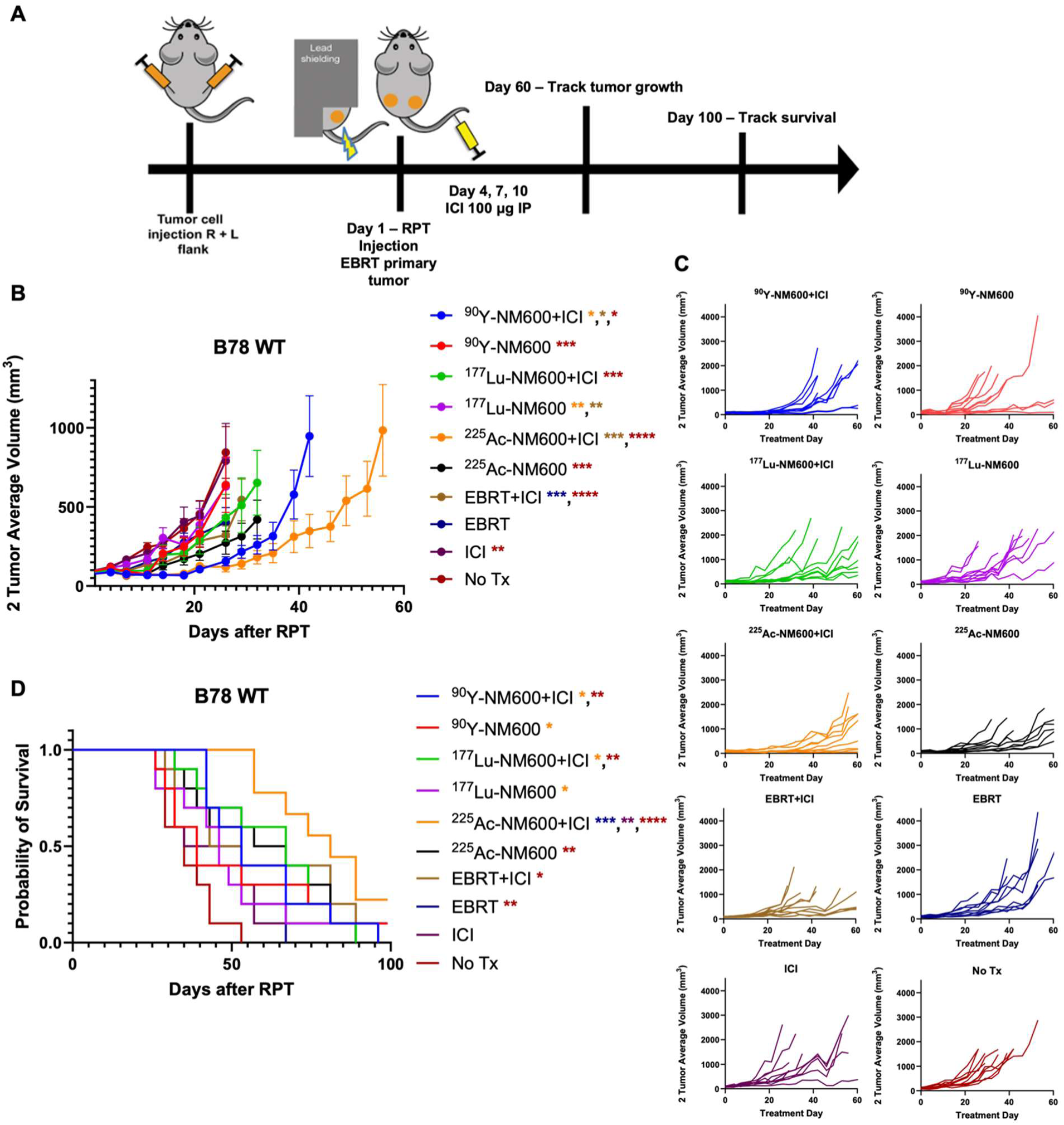
^225^Ac-NM600+ICI extends greater survival benefit than ^177^Lu-, ^90^Y-NM600+ICI in two-tumor B78 melanoma model. **(A)** Treatment scheme for *in vivo* therapy studies. **(B-D)** B78 wild-type (WT) two-tumor bearing mice were randomized to untreated control (No Tx), dual anti-CTLA4 and anti-PD-L1 (ICI), 4 Gy ^90^Y-NM600 (3.145 MBq), 4 Gy ^177^Lu-NM600 (7.104 MBq), 2 Gy ^225^Ac-NM600 (18.5 kBq), 4 Gy primary tumor-targeted EBRT, or combination ^90^Y-, ^177^Lu-, ^225^Ac-NM600 or EBRT and ICI. Combination ^225^Ac-NM600+ICI reduces tumor growth rate in B78 WT tumors compared to ^90^Y-NM600+ICI, ^177^Lu-NM600, EBRT+ICI, and No Tx groups **(B-C)** and prolongs survival in B78 WT tumor-bearing mice compared to ^90^Y-NM600+ICI, ^90^Y-NM600, ^177^Lu-NM600+ICI, ^177^Lu-NM600, EBRT, ICI, and No Tx groups **(D)**. ^225^Ac-NM600+ICI treatment group: n=9, all other treatment groups: n=10. Log-rank test was used to compare survival.

Additionally, combination ^225^Ac-NM600+ICI prolongs survival in B78 WT tumor-bearing mice compared to ^90^Y-NM600+ICI (median survival: 53 days), ^90^Y-NM600 (39 days), ^177^Lu-NM600+ICI (67 days), ^177^Lu-NM600 (46 days), EBRT (53 days), ICI (44 days), and No Tx (35 days) groups (**Figure 7D**, **Table S8**).

## Discussion

Using *in vitro* dosimetry, we established that radiation of different particle emissions leads to IFN1 responses with differing magnitudes and time courses. These results are consistent with an impact of radioisotope physical properties on immune response, with high LET, long half-life ^225^Ac leading to IFN1 responses of the greatest magnitude and longest duration compared to β-emitting ^90^Y-or ^177^Lu-NM600 *in vitro*. Radioisotope physical properties, including dose, half-life, and LET influenced expression of other immune susceptibility markers including *Fas* and *Pdl1*.

*In vivo*, we found tumor CD8^+^/Treg ratios increased following RPT and that the combined therapeutic efficacy of ^225^Ac-NM600+dual ICIs in B78 melanoma requires STING. STING KO in the B78 model suggests that the *Ifnb1* response may be earlier and less durable in the absence of STING, which may explain the loss of effect of combined ^225^Ac-NM600+dual ICI when compared to ^225^Ac-NM600 alone, as STING-independent IFN1 response may have been insufficient to promote CD8^+^ T cell infiltration.

These studies demonstrate a critical impact of radionuclide selection on the timing of TME immunomodulation by RPT, which may be important to optimizing study design and interpreting results of RPT + ICI combinations. Indeed, timing of increase of CD8^+^/Treg ratio varied with the different radioisotopes, with ^90^Y- or ^177^Lu-NM600 yielding increases on day 7, while 12 Gy ^225^Ac-NM600 increases were delayed to day 21. The delayed effect of ^225^Ac-NM600 may be due partly to prolonged dose delivery and partly to the greater relative biological effectiveness (RBE) of its high LET radiation killing more cells (tumor, immune, stromal) in the first 1-2 half-lives (i.e., 10-20 days), when its dose rate is highest. Thus, existing and newly infiltrating immune cells that border cells of ^225^Ac-NM600 uptake may be killed, delaying immunomodulation until after 1-2 half-lives when TILs are able to repopulate.

The observation of increased CD8^+^/Treg ratio in the tumor following ^90^Y-, ^177^Lu-, and ^225^Ac-NM600 only, but not EBRT or WMRT, highlights a potential advantage of RPT in sparing irradiation of secondary lymphoid organs, as surgical removal (*41*) or elective irradiation (*42*) of tumor draining lymph nodes has been shown to mitigate response to radiotherapy. The negative results of clinical trials combining radiotherapy and immunotherapy (*43, 44*) may in part be explained by inclusion of lymph nodes in the radiation field, thus killing antigen presenting and effector cells necessary for systemic response. RPT, particularly short-range agents such as ^225^Ac-NM600, may be able to overcome this barrier when successfully targeted against tumor-specific molecules or antigens, sparing damaging radiation to neighboring lymphoid structures.

Additionally, the greater therapeutic efficacy of ^225^Ac-NM600+ICI over ^90^Y-NM600 (tumor growth, survival), ^177^Lu-NM600+ICI (survival), and EBRT+ICI (tumor growth) in a two-tumor immunologically cold tumor model suggests that an alpha particle-emitting, short range, high LET RPT agent may yield the greatest clinical benefit in combination with ICI compared to other RT modalities.

These findings suggest capitalizing on immunomodulatory effects from ^225^Ac-NM600 and other alpha-particle RPT agents may require: 1) administering ICI before or after RPT (non-concurrently), 2) combining long half-life RPT agents with ICI agents with long circulation times, or 3) combining tumor-targeting RPT with non-tumor-targeting immunotherapies (i.e., immune organ-targeting therapies). A combination of these strategies may ensure that the intended immune effects at the tumor are not abrogated by concurrent high LET emissions at the tumor site.

These studies had several limitations. First, one cannot isolate and study a single physical property of a radioisotope in a controlled manner. To address this, we have administered EBRT innovatively to specifically test hypotheses about dose rate and absorbed dose accumulation.

Nonetheless, with radioisotope studies we cannot rule out that differences arise from multiple chemical and physical properties, and we report correlations here for immune response to various radioisotopes without being able to definitively conclude about the specific effects of individual physical properties. A more controlled study to experimentally isolate and test the impact of a radioisotope physical property on immune responses may not be feasible. Our data suggest that observed differences in immune response correlate with physical properties of radioisotopes. The data do not prove this correlation, but they are consistent with it.

Additionally, we acknowledge that the absorbed doses determined by our dosimetry methods are estimates, and factors not included in our models, particularly the assumption that all daughter isotopes of actinium are not redistributed, may alter the estimates of absorbed dose presented here. In vitro dosimetry was unable to be determined for the Auger-emitting radionuclides with our model; therefore, we do not have estimates for how the activities in those experiments may compare to the in vitro ^90^Y, ^177^Lu, and ^225^Ac doses.

Another constraint is contact inhibition of cells in control groups, which limits *in vitro* assays to 7 days of irradiation. Additionally, the low, 4 Gy dose used in B78 experiments prevented optimal evaluation of IFN1and other immunomodulatory responses following EBRT, ^90^Y, and ^177^Lu; however, it did highlight the unique potential of ^225^Ac to activate IFN1 responses at low dose. *In vivo*, although syngeneic murine tumor models may be limited in their ability to model human tumor heterogeneity and immune contexture, they can provide insight into how the different radiobiological properties of certain RPT agents may affect tumor immunomodulation timing, magnitude, and location compared to observations made following EBRT. Given the difficulty of imaging ^225^Ac with the low activities (9.25-18.5 kBq) used in these preclinical studies, ^225^Ac-NM600 dosimetry was limited to time-activity curves calculated from *ex vivo* biodistribution data. However, the potential to perform patient-specific SPECT-based dosimetry for ^225^Ac compounds is of great interest to the medical physics community, and the feasibility of doing so has been reported (*45*).

Lastly, we and others have previously reported that TIL populations show the most significant change following immunotherapy or combination of radiation and immunotherapy, as opposed to radiation monotherapy (*6, 16*). Surprisingly, we demonstrate here that RPT alone modulates immune populations in a tumor-specific manner, perhaps related to tumor-draining lymph node preservation following short-range RPT. While it is important to understand this independent effect of RPT, it is also critical to evaluate immune cell population changes following combination radiation therapy and immunotherapies to understanding mechanisms of response and resistance. Additional studies evaluating lymphocyte activation and a more detailed profile of myeloid populations in future studies are warranted to test the hypothesis that RPT stimulates an activated TME while also preserving antigen presentation at the tumor-draining lymph node. Understanding RPT effects on the TME may enable rational design of clinical studies that integrate RPT and immunotherapies into patient care to enhance anti-tumor immunity in the setting of metastatic cancers.

The key observations from this study are that at low to moderate doses, ^90^Y, ^177^Lu, and ^225^Ac-based radiopharmaceuticals immunomodulate the tumor microenvironment, including type I interferon response activation in tumor cells and altering tumor infiltrating lymphocyte composition. Our data suggest that the timing and magnitude of type I interferon response gene expression following RPT treatment delivering ^90^Y, ^177^Lu, or ^225^Ac are potentially influenced by the half-life and LET of each radionuclide. ^225^Ac-NM600 elicits cooperative therapeutic efficacy in combination with ICIs to achieve STING-dependent anti-tumor responses in an immunologically cold two-tumor mouse model.

## Materials and Methods

### Cell lines and culture

The murine head and neck cancer MOC2 cell line was generously provided by Dr. Ravindra Uppaluri. The murine melanoma B78-D14 (B78) cell line, derived from B16 melanoma as previously described, was obtained from Dr. Ralph Reisfeld (Scripps Research Institute) in 2002 (*46*). *Tmem173* -/-CRISPR deletion B78 (STING KO) melanoma was generated by our lab, as previously described (*47*). Cell line authentication was performed per ATCC guidelines using morphology, growth curves, and Mycoplasma testing within 6 months of use. Wild-type murine melanoma B78 (B78 WT), B78 STING KO, and MOC2 cells were grown in a humidified incubator at 37°C with 5% CO_2_, in RPMI-1640 supplemented with 10% FBS, 100 U/mL penicillin, and 100 µg/mL streptomycin. EBRT or radionuclide therapy was delivered at 60-70% cell confluency and cell media was exchanged with fresh, pre-warmed media every other day until harvest of adherent cells for analysis.

### Murine tumor models

Mice were housed and treated under a protocol approved by the Institutional Animal Care and Use Committee at the University of Wisconsin-Madison (protocol: M005670). Female C57BL/6 mice were purchased at age 6 to 8 weeks from Taconic. MOC2 tumors were engrafted by subcutaneous flank injection of 1 × 10^6^ tumor cells. B78 WT and STING KO tumors were engrafted by subcutaneous flank injection of 2 x 10^6^ tumor cells. Tumor size was determined using calipers and volume approximated as (width^2^ × length)/2. Mice were randomized immediately before treatment by permuted block randomization. Only mice with palpable flank tumors the day before treatment began were included in the study. Treatment began when tumors were well-established, approximately 2 weeks after tumor implantation for MOC2 (∼200 mm^3^) and 4 weeks for B78 WT and STING KO (∼50-100 mm^3^). The day of EBRT or RPT was defined as “day 1” of treatment. Anti-murine CTLA-4 (IgG2c, clone 9D9, produced by NeoClone) and anti-murine PD-L1 (IgG2b, clone 10F.9G2, BioXCell) were administered by 100 μg intraperitoneal injection on days 4, 7, and 10. Mice were euthanized when tumor size exceeded 20 mm in longest dimension or recommended by an independent animal health monitor for morbidity or moribund behavior. Due to institutional radiation safety protocols and the mice being treated with radioactive isotopes, all investigators were aware of mouse treatment groups.

The number of mice in therapy groups was based on anticipated effect size of outcomes to power experiments, determined in collaboration with biostatisticians, and therefore may vary between treatment groups and tumor models. Mouse therapy experiments were repeated in duplicate. Final replicates are presented for tumor response and aggregate data for survival; number of animals per group is indicated.

### Radionuclide production

#### Cobalt-58m production and separation

Cobalt-58m was produced by irradiating isotopically enriched ^57^Fe targets with 8.2 MeV deuterons using the GE PETtrace at University of Wisconsin-Madison. Targets were made and processed by cation exchange chromatography and extraction chromatography as previously described (*48*), using mixed inorganic acid/acetone solvents to isolate ^58m^Co. The purified product was evaporated to dryness and reconstituted in PBS for *in vitro* cell studies.

#### Antimony-119 production and isolation

Metallic tin targets were electrodeposited from isotopically enriched ^119^Sn (96.3% ^119^Sn, 2.6% ^118^Sn, 1.1% ^120^Sn, Isoflex) with lineal thicknesses of 100 mg/cm^2^ as reported previously (*49*). The targets were irradiated with 35 µA of degraded, 12.6 MeV proton beam for 1 h using a GE PETtrace cyclotron. Irradiated targets were dissolved in 2 mL concentrated HCl at 90 °C for 1h. A 10x dilution of the dissolved target solution was loaded onto 100 mg mercaptopropyl functionalized silica gel resin (Sigma-Aldrich, 200-400 mesh) to isolate no-carrier-added ^119^Sb from bulk ^119^Sn target material (*50*). The resin was washed with 0.5 mL acidic ethanol solution prepared by combining 0.1 mL 10.5M HCl with 10.4 mL EtOH. Antimony was eluted from the resin using 0.75 mL acidic ethanol solution prepared by combining 5 mL 10.5M HCl with 5.5 mL EtOH. The acidic ethanol elution was dried to a small volume (<50 μL) using N_2_ gas and diluted with 10 mL PBS.

### Radiochemistry

^90^Y was purchased as ^90^YCl_3_ from Eckert and Ziegler. ^177^Lu and ^225^Ac were purchased as ^177^LuCl_3_ and solid ^225^Ac(NO_3_)_3_ from Oak Ridge National Laboratory. The radiolabeling of 2- [[hydroxy[[18-[4-[[2-[4,7,10-tris(carboxymethyl)-1,4,7,10-tetraazacyclododec-1-yl]acetyl]amino]phenyl]octadecyl]oxy]phosphinyl]oxy]-N,N,N-trimethyl ethanaminium (NM600) with ^90^Y, ^177^Lu, or ^225^Ac proceeded by mixing the radiometal with NM600 (0.336-3.36 MBq/nmol) in 0.1 M NaOAc buffer (pH 5.5) for 30 min at 90°C, as previously described for ^90^Y-NM600 (*34*). The labeled compounds were purified using a reverse-phase Waters Oasis HLB Light cartridge (Milford, MA), eluted using 100% ethanol, dried with a stream of N_2_, and reconstituted in 0.9% NaCl with 0.1% v/v Tween 20. Yield and purity were determined with instant thin-layer chromatography using 50 mM EDTA and iTLC-SG –glass microfiber chromatography paper impregnated with silica gel (Agilent Technologies). A cyclone phosphor image reader was used to analyze the chromatograms; the labeled compound remained at the spotting point while free radiometals moved with the solvent front.

### SPECT/CT imaging

Animals with subcutaneous MOC2 or B78 WT (C57BL/6) tumors were administered 18.5 MBq of ^177^Lu-NM600 in the lateral tail vein. Individual mice (n=3) were placed prone into a MILabs U-SPECT6 /CTUhr system (Houten, The Netherlands) under 2% isoflurane for longitudinal scans at 3, 24, 72, 168, and 240 h post-injection (240 h: MOC2 only). CT scans (5 min) were acquired for anatomical reference and attenuation correction and fused with the SPECT scans (45 min). Image reconstruction used a similarity-regulated ordered-subset expectation maximization (SROSEM) algorithm. Images were quantitatively analyzed by drawing volumes of interest (VOI) over the tumor and organs of interest to determine the percent injected activity (IA) per gram (%IA/g) for each tissue. An *ex vivo* biodistribution study was carried out after the last scan time point.

### *Ex vivo* biodistribution

Mice bearing MOC2, B78 WT, or B78 STING KO (^225^Ac-NM600 only) subcutaneous tumors were injected with 18.5 MBq of ^177^Lu-NM600 or 9.25 kBq of ^225^Ac-NM600, then euthanized via CO_2_ asphyxiation at 3, 24, 48, 72, 216, and 384 h (n = 3) post-injection, and organs of interest were collected for *ex vivo* biodistribution. Organs from animals that received ^225^Ac-NM600 were allowed to decay at 4°C overnight to reach secular equilibrium with ^213^Bi, which could be quantified. Organs were wet weighed, analyzed with a Perkin Elmer Wizard2 Gamma Counter (Westham, MA), and decay corrected to calculate the %IA/g for each tissue.

### *In vitro* dosimetry estimation and validation

Cells were irradiated with cell culture media containing specific activities of ^90^Y, ^177^Lu, or ^225^Ac estimated by GEANT4 Monte Carlo to deliver 12 Gy (MOC2) and 4 Gy (B78) to a cell monolayer at an infinite timepoint in a 6-well plate. Specific activities added for each dose of each radioisotope are listed in **Table S1**. A stock solution of radionuclide-containing cell culture media was prepared to maintain specific activity throughout media changes (**Figure S1**). To validate the Monte Carlo dosimetry predictions, a separate experiment was performed mimicking the cell irradiation scenario to measure the absorbed dose with thermoluminescent dosimeters (TLDs) and Gafchromic film. TLDs were placed in the center of wells of a 6-well cell culture plate to evaluate the absolute absorbed dose and Gafchromic film was placed underneath the cell culture plate to evaluate the homogeneity of the absorbed dose throughout the well. A serial dilution of low- to moderate-dose free ^90^Y activity concentrations was added to the wells and exposed the TLDs and film. After 10 days of exposure, the TLDs were collected and analyzed by the University of Wisconsin-Madison Radiation Calibration Laboratory (Calibration Cert #1664.01). Using the TLDs we had available and for which we have experience, it was not feasible to measure the absorbed dose of ^177^Lu and ^225^Ac because of each isotope’s emission radiological pathlength and the subsequent partial volume effects, which would not be accounted for in the TLD calibration process. As such, our emphasis in this paper was to move past a computational study alone and validate our computational model with the ^90^Y predictions. This validation imparted confidence to simulations conducted with subsequent isotopes. The ^90^Y validated Monte Carlo dosimetry tool was then updated with different isotopes, radioactive decay constants and half-lives for ^177^Lu and ^225^Ac estimates.

### *In vivo* dosimetry estimation

To estimate dosimetry of ^225^Ac-NM600, *ex vivo* biodistribution results were used. Allometric scaling was first performed to estimate the organ mass with respect to body mass, assuming 20 g for C57BL/6. Then, the residence time in each organ, MBq-sec/MBq(inj), was determined using trapezoidal integration, assuming only physical decay after the last timepoint. Lastly, the residence time for each organ was multiplied by the dose factor, mGy/MBq-sec, for each organ, which was assumed to be a sphere of equal mass that only receives self-dose. The total dose for each organ was computed by summing all contributions from the complete decay chain of ^225^Ac, considering negligible redistribution of daughter isotopes. SPECT/CT-based ^177^Lu-NM600 dosimetry was estimated according to a previously described method (*35, 36*).

Total injected activity within each organ was calculated from extrapolation of %IA/g at a given time point. The standard mouse model was used to convert organ-specific cumulative activity into absorbed dose per injected activity (Gy/MBq). Dose contributions from surrounding organs were also included in calculations.

### Radiation therapy (RT)

Delivery of EBRT (195 kV) *in vitro* was performed using a RS225 Cell Irradiator (Xstrahl). EBRT regimens were 12 Gy for MOC2 or 4 Gy for B78, as well as regimens designed to mimic radionuclide (^90^Y, ^177^Lu, ^225^Ac) delivery of the same doses, as determined by exponential decay (Dose delivered in time ***t*** = Total Dose*(0.5)***^t^***^/**t**^_1/2_). For “dose equivalent” EBRT, cells were irradiated on day 0 with the dose delivered by each radionuclide by each cell harvest timepoint (days 1, 3, 7). For “dose fractionation” EBRT, cells were irradiated every 24h with the dose delivered by each radionuclide in that interval. All radiation doses for each EBRT regimen are listed in **Table S2**.

^90^Y-, ^177^Lu-, or ^225^Ac-NM600 treatment (at activities of 3.7 MBq, 7.4 MBq, or 9.25 kBq, respectively, per mouse) to achieve a tumor dose prescription of 12 Gy was administered via intravenous tail vein injection on treatment day 1 for MOC2 tumor-bearing mice. For B78 WT or B78 STING KO tumor-bearing mice, 3.15 MBq ^90^Y-NM600, 7.10 MBq ^177^Lu-NM600, or 18.5 kBq ^225^Ac-NM600 per mouse was administered via intravenous tail vein injection on treatment day 1 to achieve a tumor dose prescription of 4 Gy for ^90^Y- and ^177^Lu-NM600 and 2 Gy for ^225^Ac-NM600. Delivery of EBRT (300 kV) *in vivo* was performed using an X-ray biological cabinet irradiator X-RAD CIX3 (Xstrahl). The dose rate for EBRT delivery in all experiments was approximately 2 Gy/min. Dosimetric calibration and monthly quality assurance checks were performed on these irradiators by University of Wisconsin Medical Physics staff. Tumor-targeted EBRT was delivered at a dose of 12 Gy (MOC2) or 4 Gy (B78) on day 1 with the tumor exposed and the rest of the animal shielded using custom lead blocks. For whole-mouse radiation, animals were placed in a custom clear plastic restraint and treated posterior to anterior to a calculated midline dose of 12 Gy without any shielding.

### Gene expression analysis

Cells treated *in vitro* with EBRT or ^90^Y, ^177^Lu, or ^225^Ac were washed with cold PBS. TRIzol reagent (ThermoFisher Scientific Cat #15596026) was added to the plate, and the cells were collected via scraping. For analysis of tumor tissue, tumors were harvested, and samples were homogenized in TRIzol using a Bead Mill Homogenizer (Bead Ruptor Elite, Omni International Cat #19-040E). For *in vitro* and *in vivo* samples, total RNA was extracted using RNeasy Mini Kit (QIAGEN, Germany, Cat #74106) according to the manufacturer’s instructions. Extracted RNA was subjected to complementary cDNA synthesis using QuantiTect Reverse Transcription Kit (QIAGEN, Germany, Cat #205314) according to the manufacturer’s instructions. Quantitative PCR (qRT-PCR) was performed using TaqMan Fast Advanced qPCR Master Mix (ThermoFisher Scientific Cat #4444556). Thermal cycling conditions (QuantStudio 6, Applied Biosystems) included the UDG activation at 50°C for 2 min, followed by Dual-Lock DNA polymerase activation stage at 95°C for 2 min followed by 40 cycles of each PCR step (denaturation) 95°C for 1 s and (annealing/extension) 60°C for 20 s. For data analysis, the Ct values were exported to an Excel file, and fold change normalized to untreated control samples was calculated using the ΔΔCt method. *Hprt* was used as an endogenous control. A complete list of TaqMan probes is included in **Table S3**. *In vitro* qPCR experiments were repeated in duplicate (EBRT-eq, EBRT-fr, ^58m^Co) or triplicate (no treatment, EBRT, ^90^Y, ^177^Lu, ^225^Ac); final replicates are presented. Number of wells or animals per group is indicated.

### In vitro immunofluorescence

Before radiation, cells were seeded onto poly-L-lysine-coated coverslips in 6-well Corning Costar^®^ TC-treated plates (Cat #3516). Warmed RPMI cell growth medium was applied at 3 mL per well. MOC2 and B78 cells were treated with EBRT or radionuclide-containing media, as described above. At the fixation timepoint (1 h; 1, 3, or 7 days), the cells were washed with PBS, fixed with 4% paraformaldehyde solution, then stored in PBS at 4°C. At the end of the seven-day time course (EBRT, control) or when the cells had reached background levels of radiation (radionuclide therapy), cells were permeabilized with 0.1% Triton X-100 and non-specific binding was blocked with SuperBlock (TBS T20, ThermoFisher). The primary antibody (Phospho-Histone H2A.X (Ser139) (20E3) Rabbit mAb, #9718, Cell Signaling Technology, 1:400) was applied and incubated overnight at 4°C. After washing, an AlexaFluor 488 labeled secondary antibody (ThermoFisher, Catalog #A-11008, 1:1000) was applied and incubated for 1 h at room temperature; the cell nuclei were stained with DAPI. The coverslips were mounted with ProLong Gold Antifade Mountant (ThermoFisher, Cat #P36930). 3-5 representative fluorescent images per sample were taken using the 60x/1.4 NA oil objective on a Nikon A1R-s LSCM Confocal Microscope and processed using Nikon NIS-Elements viewer. Using ImageJ for image analysis, γH2AX foci were manually quantified for 50 nuclei/sample.

### Flow cytometry

Flow cytometry was performed as previously described (*51*), using fluorescent beads (UltraComp Beads eBeads, 176 Invitrogen) to determine compensation, and fluorescence minus one (FMO) methodology to determine gating. Rainbow beads (Spherotech) were used to set voltages for each flow cytometry timepoint. For *in vivo* analysis, tumors were harvested and gently dissociated. Spleens and tumor draining lymph nodes were harvested and manually dissociated. Bone marrow was flushed from the tibia with phosphate-buffered saline and strained. Blood was collected by submandibular vein collection. Bone marrow, spleens, and blood were treated with RBC lysis buffer (BioLegend) and washed with phosphate-buffered saline prior to staining. Cells were treated with CD16/32 antibody (BioLegend) to prevent non-specific binding. Live cell staining was performed using Ghost Red Dye 780 (Tonbo Biosciences) according to manufacturer’s instructions. After live-dead staining, a single cell suspension was labeled with the surface antibodies at 4°C for 30 min and washed three times using flow buffer (2% FBS + 2 mM EDTA in PBS). For intracellular staining, the cells were fixed and stained for internal markers with permeabilization solution according to manufacturer’s instructions (BD Cytofix/Cytoperm^TM^). Flow cytometry was performed using an Attune NxT Flow Cytometer (ThermoFisher). Data were analyzed using FlowJo Software. A complete list of antibody targets, clones, and fluorophores is provided in **Table S4**. Number of animals per group is indicated.

### Statistical analysis

Prism 10 (GraphPad Software) and R 4.1.2 (R Foundation) were used for all statistical analyses. Student’s t-test was used for two-group comparisons. Two-way ANOVA with Tukey’s honestly significant difference (HSD) test was used to assess statistical significance of observed mean differences in gene expression, flow cytometry, and *in vitro* immunofluorescence data. For tumor growth analysis, all available data was used. To compare WT and KO cell lines within each treatment group as well as the 10 treatment groups, linear mixed models after log base 10 transformation of tumor volume were fitted on cell line, time in days, and their interaction. Mouse ID was also included as a random intercept, accounting for correlation between measurements taken from the same mouse. Pairwise contrasts were adjusted using Tukey’s method (**Table S5, S7**). To compare the four treatment combinations within each cell line, linear mixed models were again used, retaining log-transformed tumor volumes and mouse ID as a random effect. The fixed effects were Ac & ICI treatment conditions, time in days, all two-way interactions, and the three-way interaction. Tukey’s HSD was used to make post-hoc pairwise comparisons of the three-way interaction (**Table S6**). Kaplan-Meier method was used to estimate the survival distribution for the overall survival. Then, pairwise comparison of the overall survival was made using a log-rank test with Benjamini-Hochberg adjustment of p-values between levels of factors (**Table S8**). All data are reported as mean ± standard error of the mean (SEM) unless otherwise noted. For all graphs, *, P < 0.05; **, P < 0.01; ***, P < 0.001; and ****, P < 0.0001.

## Acknowledgments

The authors would like to thank the University of Wisconsin Carbone Cancer Center (UWCCC), University of Wisconsin Small Animal Imaging & Radiotherapy Facility, and the UWCCC Flow Cytometry Laboratory for supporting this project. The authors would like to thank Archeus Technologies for providing NM600. The authors thank Dr. Wes Culberson and Cliff Hammer for helpful discussions and assistance with *in vitro* Gafchromic film and thermoluminescent dosimeter experiments. The authors thank Dr. Albert van der Kogel and Dr. Tracy Berg for editing and insightful review of the manuscript.

## Funding

The authors’ work is supported in part by grants from NIH NCI P50CA278595 (JPW, ZSM), NIH NCI P01CA250972 (JPW, ZSM), NIH NCI P50DE026787 (JPW, ZSM), NIH NCI P30CA014520, NIH S10OD028670-01 (JPW), NIH NCI F30CA250263 (JCJ), NIH TL1TR002375 (JCJ), NIH NCI F30CA268780 (CPK), NIH T32GM140935 (CPK, JCJ), and a UW-Madison Radiology MD-PhD Graduate Student Fellowship (CPK).

## Author contributions

Animal work and analysis: CPK, AMB, JCJ, RNS, PAC

mRNA isolation and qRT-PCR: CPK, JSK, TPTN, JCJ, MP, AGS

Flow cytometry staining and analysis: CPK, AMB, WJJ, JCJ, RNS, PAC, LMZ

Immunofluorescence staining and analysis: CPK, JSK

Cyclotron radioisotope preparation: APO, WL, EAS, JWE

RPT radiochemistry: CAF, HCR, CM, RH

Animal RPT preparation and SPECT imaging: CPK, CC, CFM, RH

SPECT analysis and dosimetry: JJG, DPA, OK, BPB

Biostatistical analysis: CPK, MH, KK

Funding, planning, and supervision of the project and data interpretation: JPW and ZSM Wrote the manuscript: CPK and ZSM

All authors read and approved the final manuscript.

## Competing interests

JJG is cofounder and Chief Innovation Officer of Voximetry, Inc. HCR provides consulting services to Archeus Technologies, which holds the license rights to NM600 related technologies. RH is a member of the scientific advisory board for Archeus Technologies. BPB is cofounder and Chief Science Officer of Voximetry, Inc. JPW is a founder and Chief Science Advisor for Archeus Technologies. ZSM. is a member of the scientific advisory board for Seneca Therapeutics, Archeus Technologies, NorthStar Medical Radioisotopes, and Cali Biomedical. RH, JPW, and ZSM are inventors on patents held by the University of Wisconsin Alumni Research Foundation related to select radiopharmaceutical therapies and the interaction of radiopharmaceutical therapies with immunotherapies. All other authors declare they have no competing interests.

## Data and materials availability

All data needed to evaluate the conclusions in the paper are present in the paper and/or the Supplementary Materials.

## Supplementary Materials

**Fig. S1.**
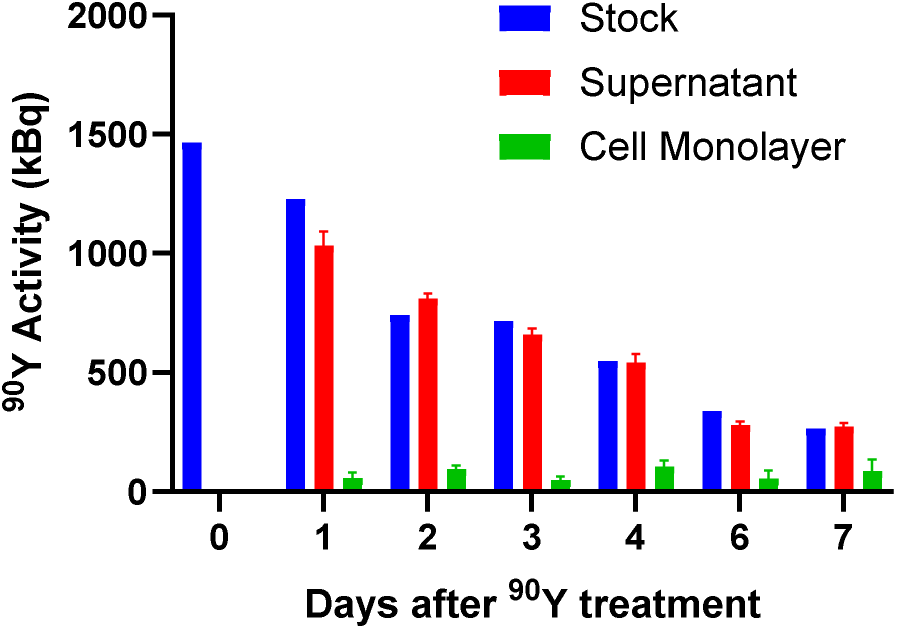
Specific activity maintained in radionuclide-containing cell culture media throughout media exchanges. 1480 kBq/mL ^90^Y in RPMI prepared as a stock solution, and ^90^Y activity in kBq measured with Capintec dose calibrator every 24h between 1-7 days after ^90^Y treatment in 1 mL of: stock solution, cell culture supernatant, or cell monolayer after washing twice with PBS and harvesting by scraping in TRIzol. N=3 for supernatant and cell monolayer, n=1 for stock solution.

**Fig. S2.**
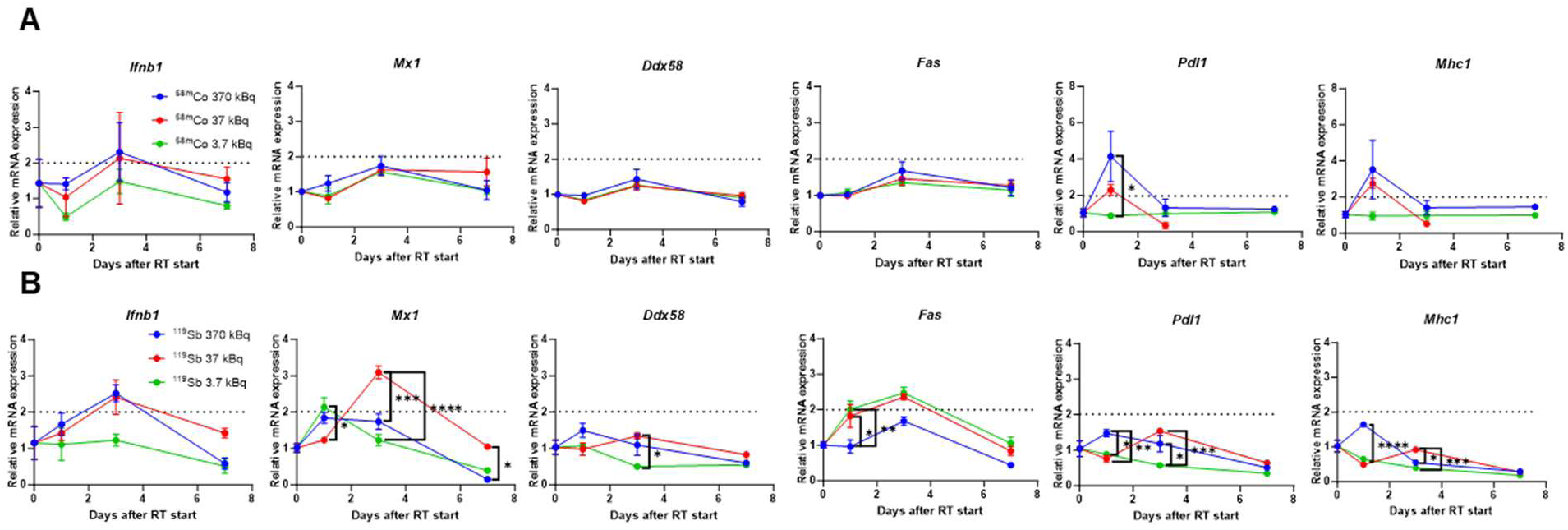
Earlier induction of type I IFN response/immunologic response to radiation following ^58m^Co and ^119^Sb. Cells were radiated with either 3.7, 37, or 370 kBq **(A)** ^58m^Co or **(B)** ^119^Sb and harvested 1, 3, or 7 days following RT. qPCR was used to quantify gene expression and is reported as fold changed normalized to untreated controls. ^58m^Co: n=3-6 per treatment group per timepoint; ^119^Sb: n=3 per treatment group per timepoint. Two-way ANOVA with Tukey’s HSD post hoc test was used to compare fold change in expression between groups.

**Fig. S3.**
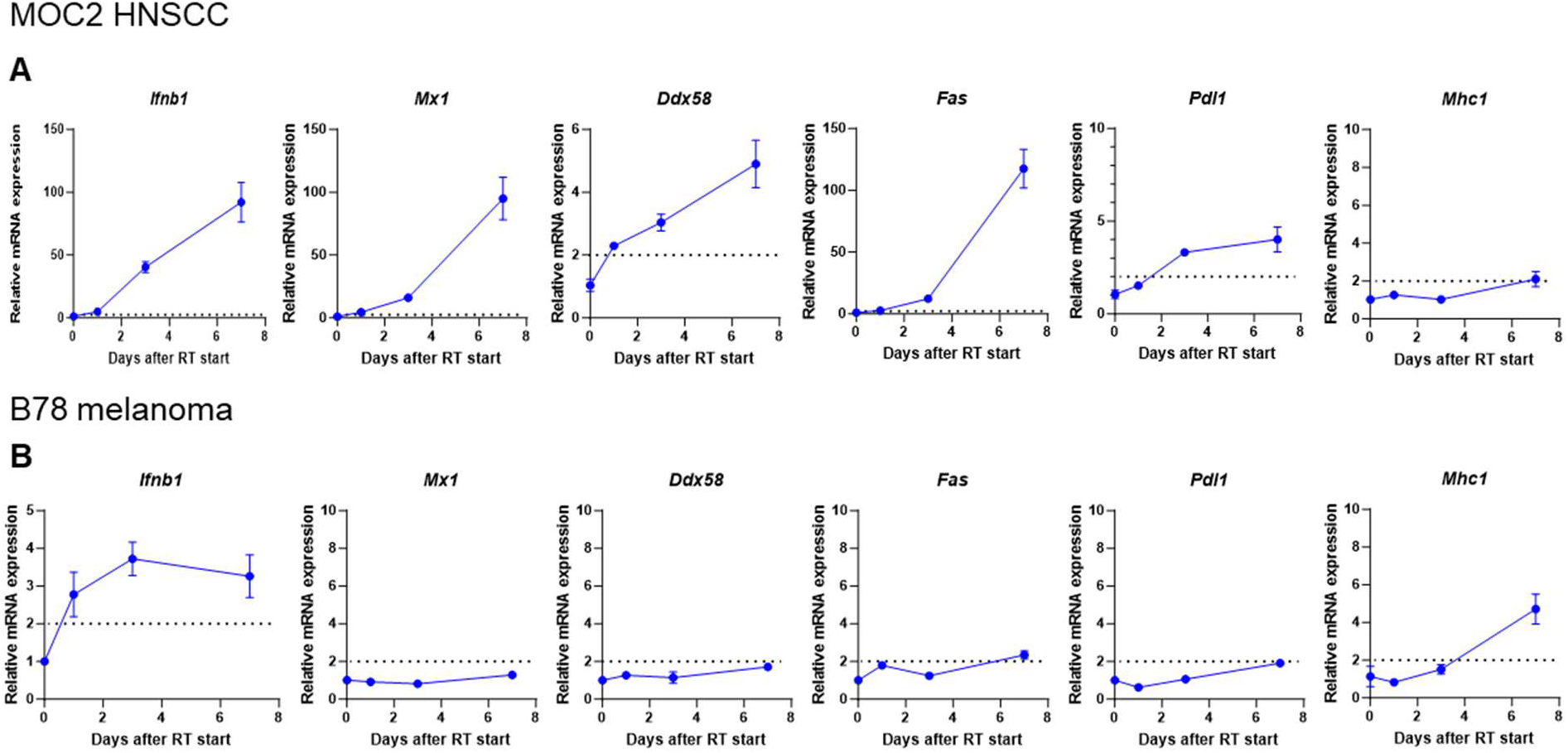
Longitudinal *Ifnb1, Mx1, Ddx58, Fas, Pdl1,* and *Mhc1* expression following (A) 12 Gy EBRT (MOC2) or (B) 4 Gy EBRT (B78) *in vitro*. Cells were harvested 1, 3, and 7 days following irradiation. qPCR was used to quantify gene expression and is reported as fold changed normalized to untreated controls. MOC2: n=6 per timepoint, B78: n=5-6 per timepoint.

**Fig. S4.**
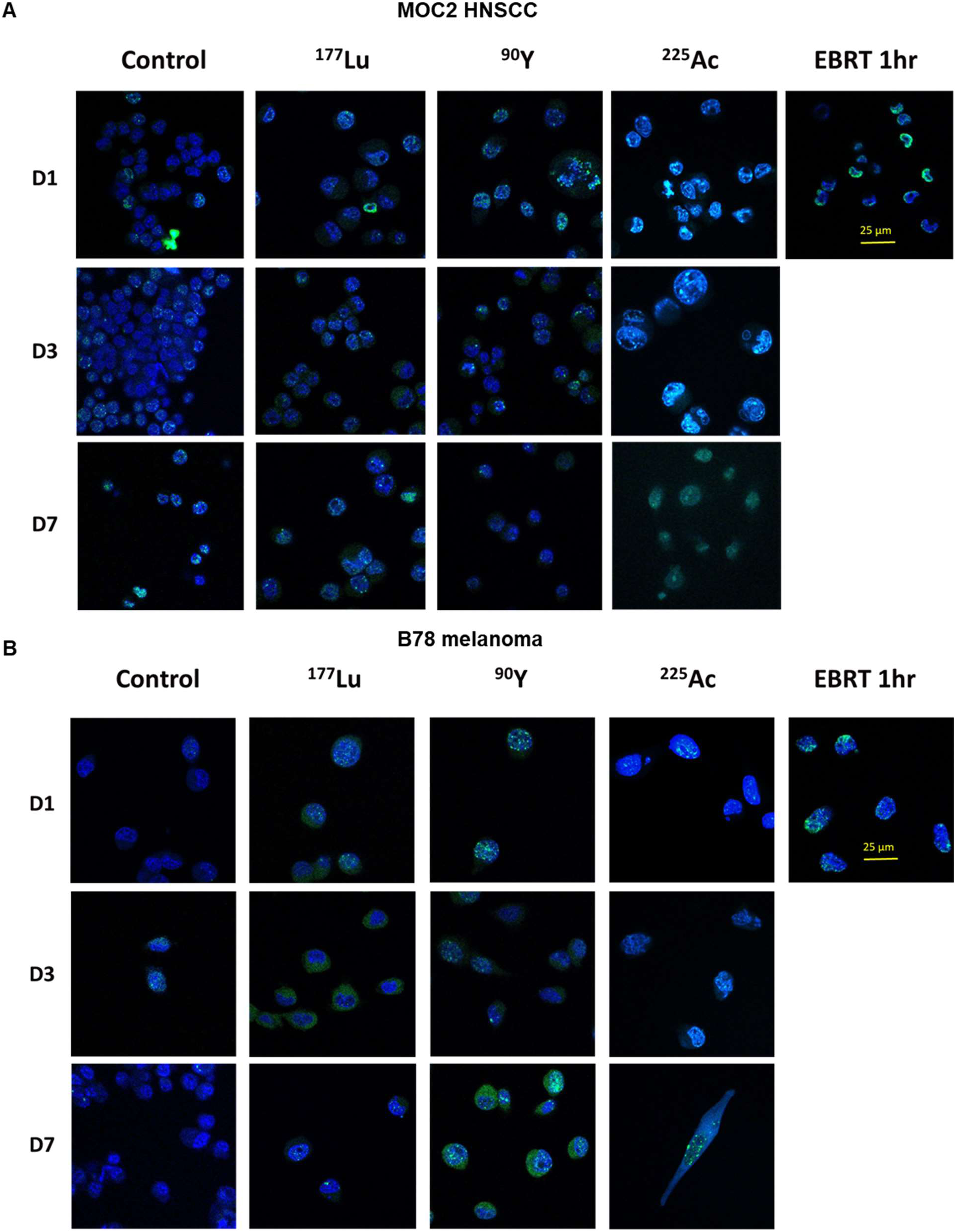
Representative images of γH2AX foci over time in (A) MOC2 (12 Gy) and (B) B78 (4 Gy) cells following EBRT, ^90^Y, ^177^Lu, and ^225^Ac. Scale bars shown. Images selected from three independent samples per treatment group; 50 cells total per treatment group quantified. γH2AX foci were not quantified for the MOC2 EBRT 1h group due to the high prevalence of apoptotic rings as opposed to nuclear foci nor for the MOC2 ^225^Ac day 7 group due to the low number of nuclei present.

**Fig. S5.**
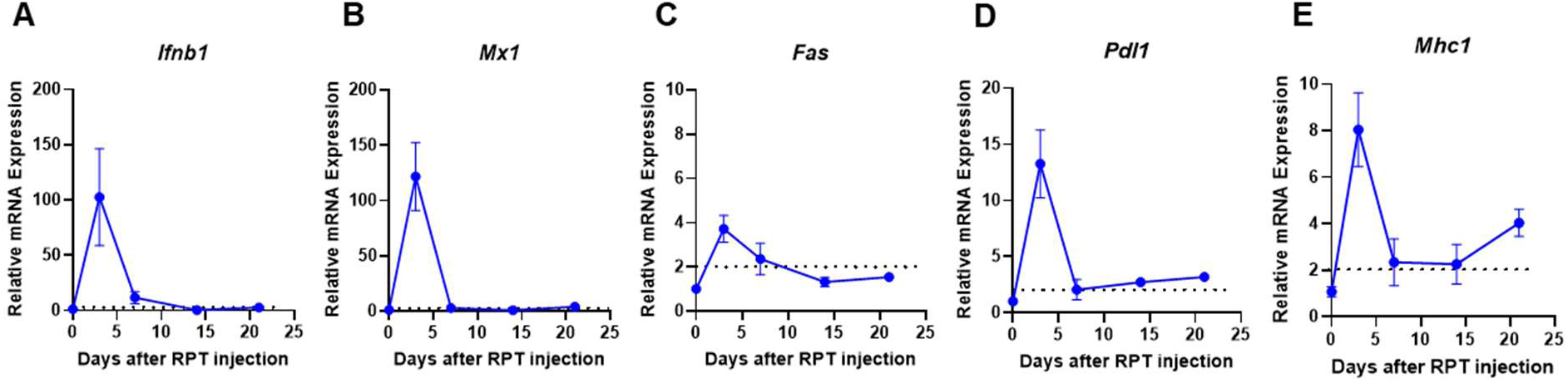
Longitudinal *in vivo* gene expression following 12 Gy EBRT (MOC2). MOC2 tumors were grown to ∼200 mm^3^ and radiated with 12 Gy tumor targeted EBRT. Tumors were harvested 3, 7, 14, or 21 days following RT administration. **(A-E)** qPCR was used to quantify gene expression and is reported as fold changed normalized to untreated controls. N=3-5 mice per treatment group per timepoint.

**Fig. S6.**
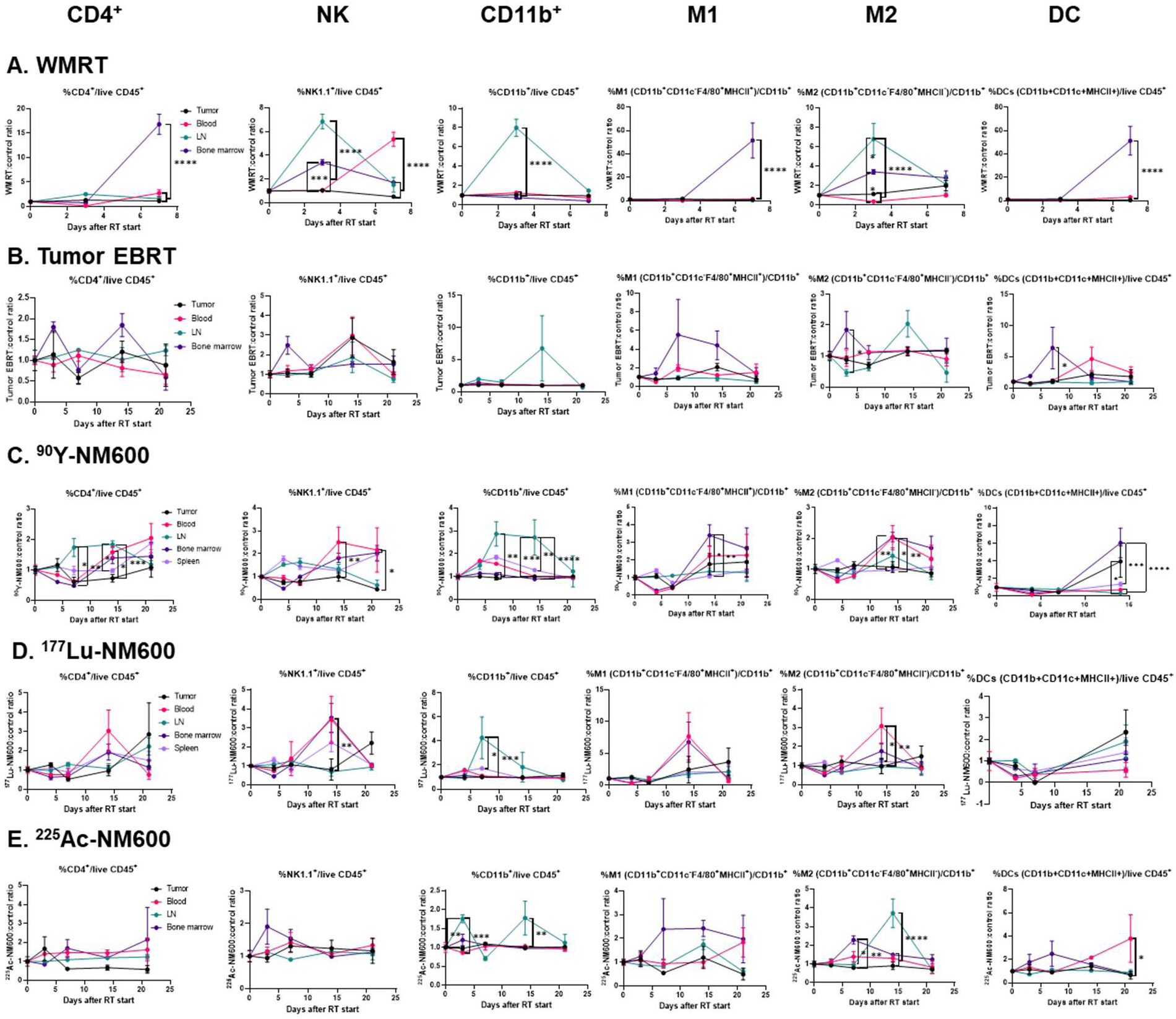
^90^Y-, ^177^Lu, ^225^Ac-NM600 modulate tumor immune cell composition in the tumor microenvironment. **(A-E)** From same mice as Figure 4, flow cytometry analyses of tumor, blood, tumor-draining lymph node, bone marrow, and spleen immune cell infiltrates [CD4^+^ T cells, NK cells (NK1.1^+^), myeloid cells (CD11b^+^), M1-like macrophages (CD11b^+^CD11c^-^F4/80^+^MHCII^+^), M2-like macrophages (CD11b^+^CD11c^-^F4/80^+^MHCII^-^), and dendritic cells (CD11b^+^CD11c^+^MHCII^+^)] as a percent of total live cells normalized to mean of control (no treatment) is shown at 3, 7, 14, and 21 days after RT administration in MOC2 HNSCC. N=4-6 per treatment group per timepoint. Two-way ANOVA with Tukey’s HSD post hoc test was used to compare ratios between tissue types.

**Fig. S7.**
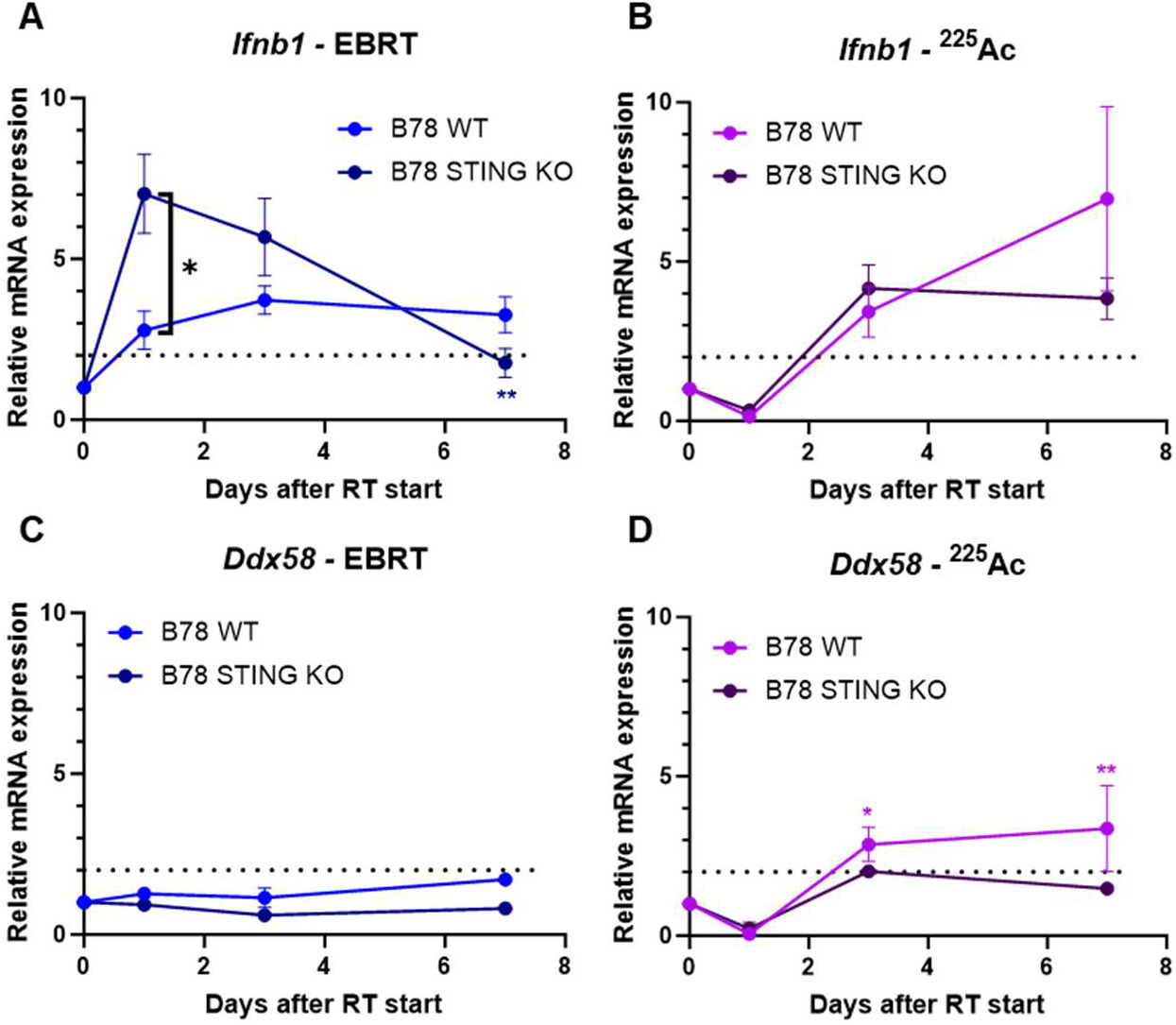
B78 WT vs. STING KO cell line in vitro *Ifnb1* and *Ddx58* gene expression upregulation following ^225^Ac or EBRT. **(A-D)** *In vitro* qPCR following 4 Gy EBRT or ^225^Ac. qPCR was used to quantify gene expression and is reported as fold changed normalized to untreated controls. qPCR: n=4-6 per treatment group per timepoint. Two-way ANOVA with Tukey’s HSD post hoc test was used to compare fold change in expression between cell lines. Colored asterisks correspond to comparisons to day 1 for EBRT (blue) or ^225^Ac (purple). Black asterisks correspond to comparisons between cell lines at the indicated timepoint.

**Table S1.**
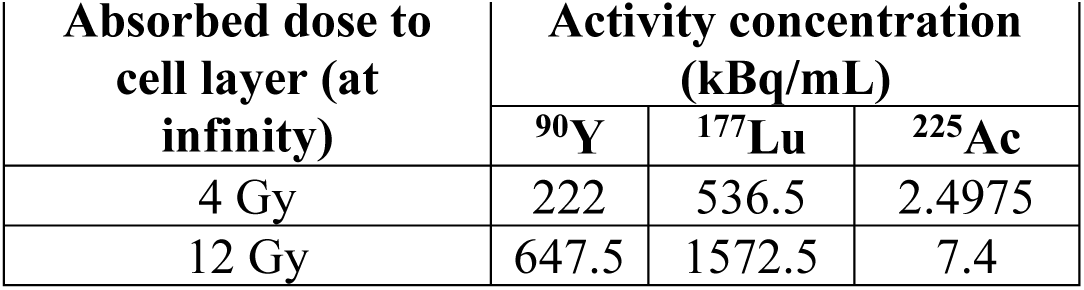
Initial radionuclide activity (kBq) required per milliliter of media as a function of mean absorbed dose (Gy) to cell monolayer at an infinite timepoint

**Table S2.**
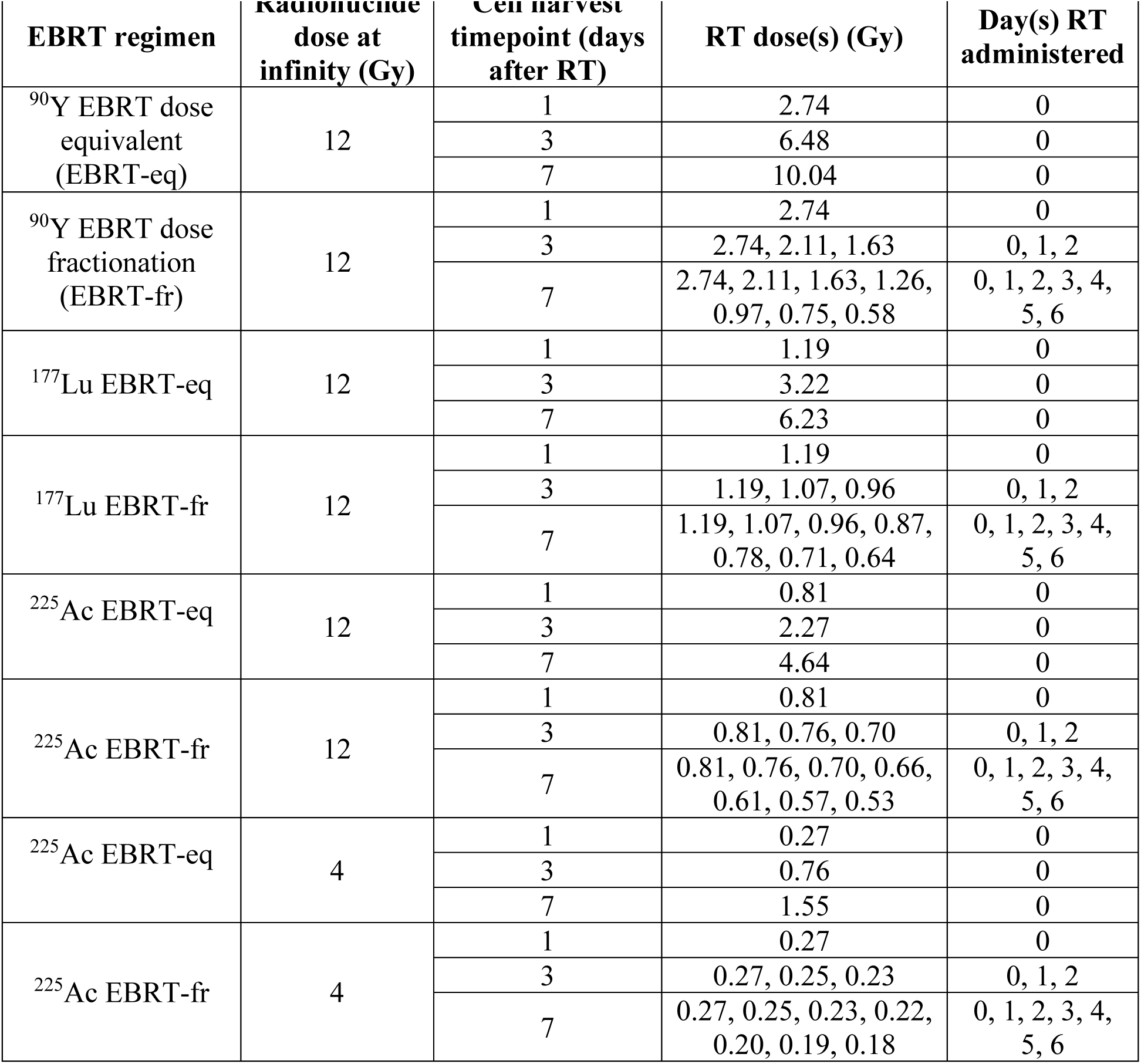
EBRT regimens for *in vitro* studies

**Table S3.**
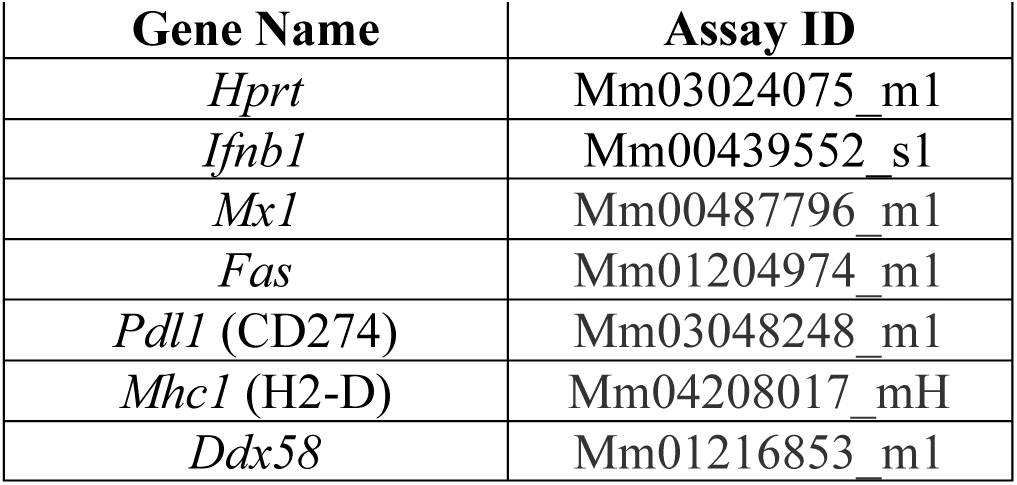
List of TaqMan probes utilized for quantitative RT-PCR experiments

**Table S4.**
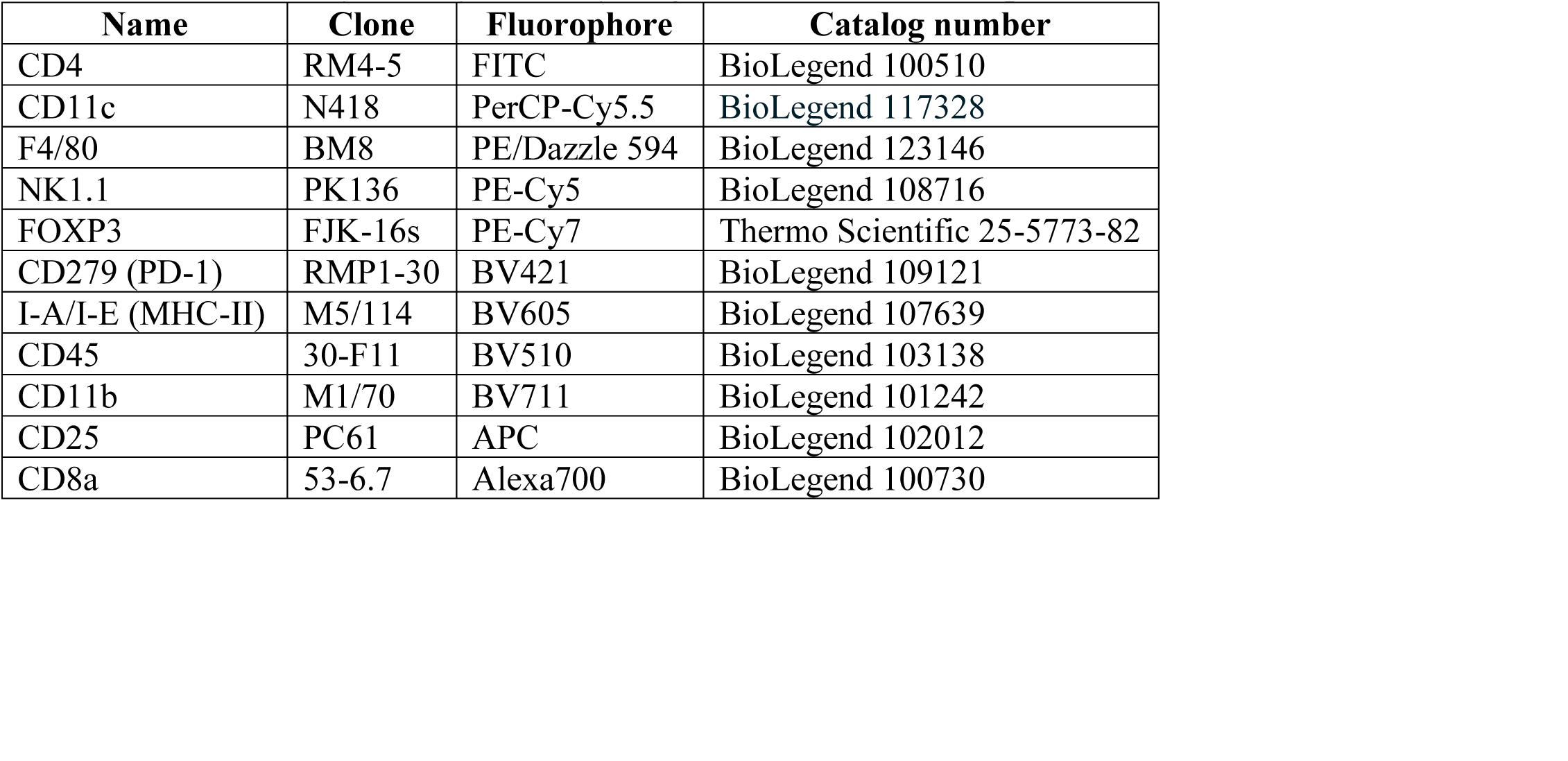
List of flow cytometry antibody targets, clones, and fluorophores

**Table S5.**
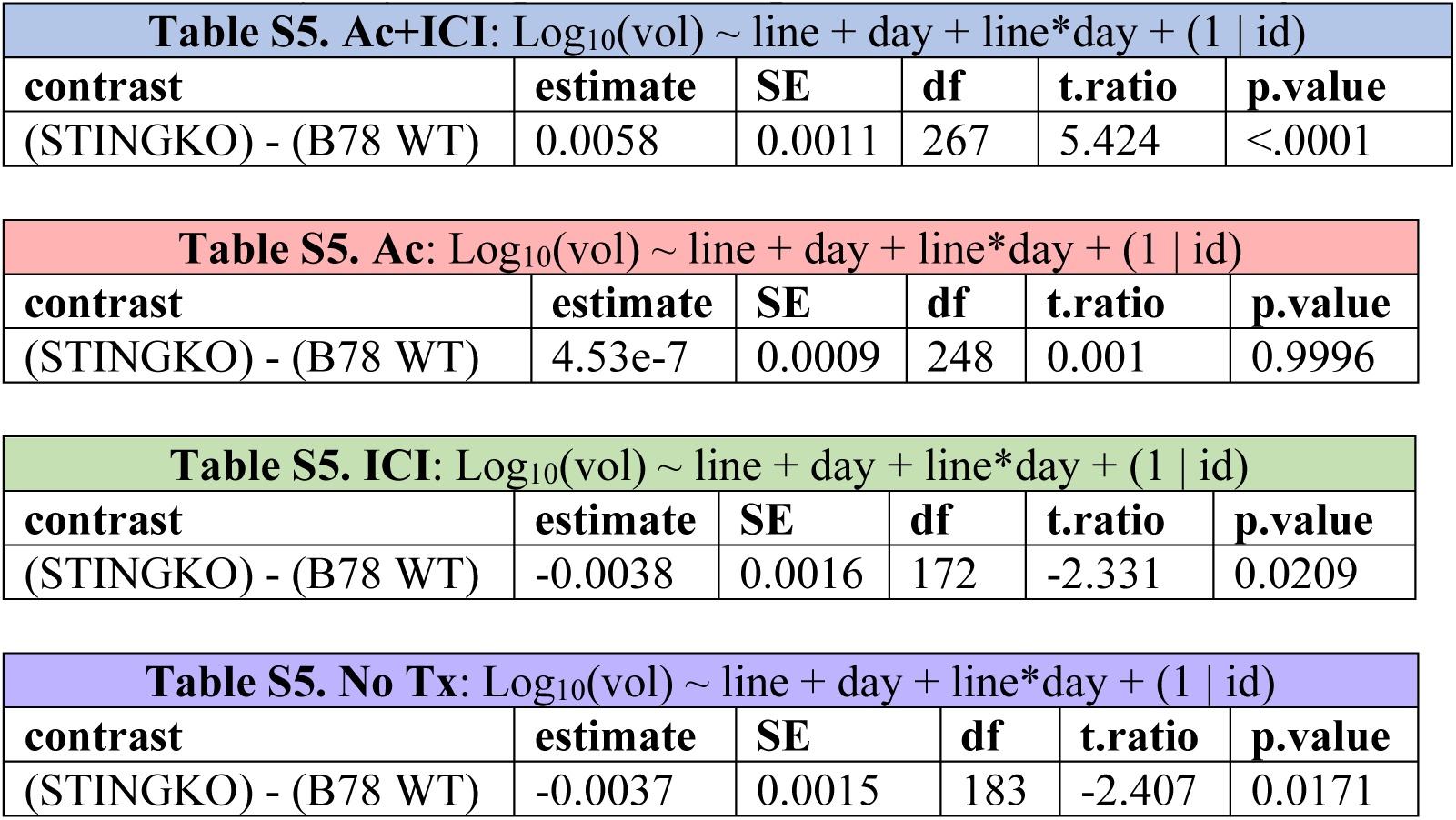
Tukey-adjusted pairwise comparisons of estimated marginal means

| **Table S5. Ac+ICI: Log<sub>10</sub>(vol) ~ line + day + line\*day + (1 \| id)** |  |  |  |  |  |
| --- | --- | --- | --- | --- | --- |
| **contrast** | **estimate** | **SE** | **df** | **t.ratio** | **p.value** |
| (STINGKO) - (B78 WT) | 0.0058 | 0.0011 | 267 | 5.424 | <.0001 |

| **Table S5. Ac: Log<sub>10</sub>(vol) ~ line + day + line\*day + (1 \| id)** |  |  |  |  |  |
| --- | --- | --- | --- | --- | --- |
| **contrast** | **estimate** | **SE** | **df** | **t.ratio** | **p.value** |
| (STINGKO) - (B78 WT) | 4.53e-7 | 0.0009 | 248 | 0.001 | 0.9996 |

| **Table S5. ICI: Log<sub>10</sub>(vol) ~ line + day + line\*day + (1 \| id)** |  |  |  |  |  |
| --- | --- | --- | --- | --- | --- |
| **contrast** | **estimate** | **SE** | **df** | **t.ratio** | **p.value** |
| (STINGKO) - (B78 WT) | -0.0038 | 0.0016 | 172 | -2.331 | 0.0209 |

| **Table S5. No Tx: Log<sub>10</sub>(vol) ~ line + day + line\*day + (1 \| id)** |  |  |  |  |  |
| --- | --- | --- | --- | --- | --- |
| **contrast** | **estimate** | **SE** | **df** | **t.ratio** | **p.value** |
| (STINGKO) - (B78 WT) | -0.0037 | 0.0015 | 183 | -2.407 | 0.0171 |

**Table S6.**
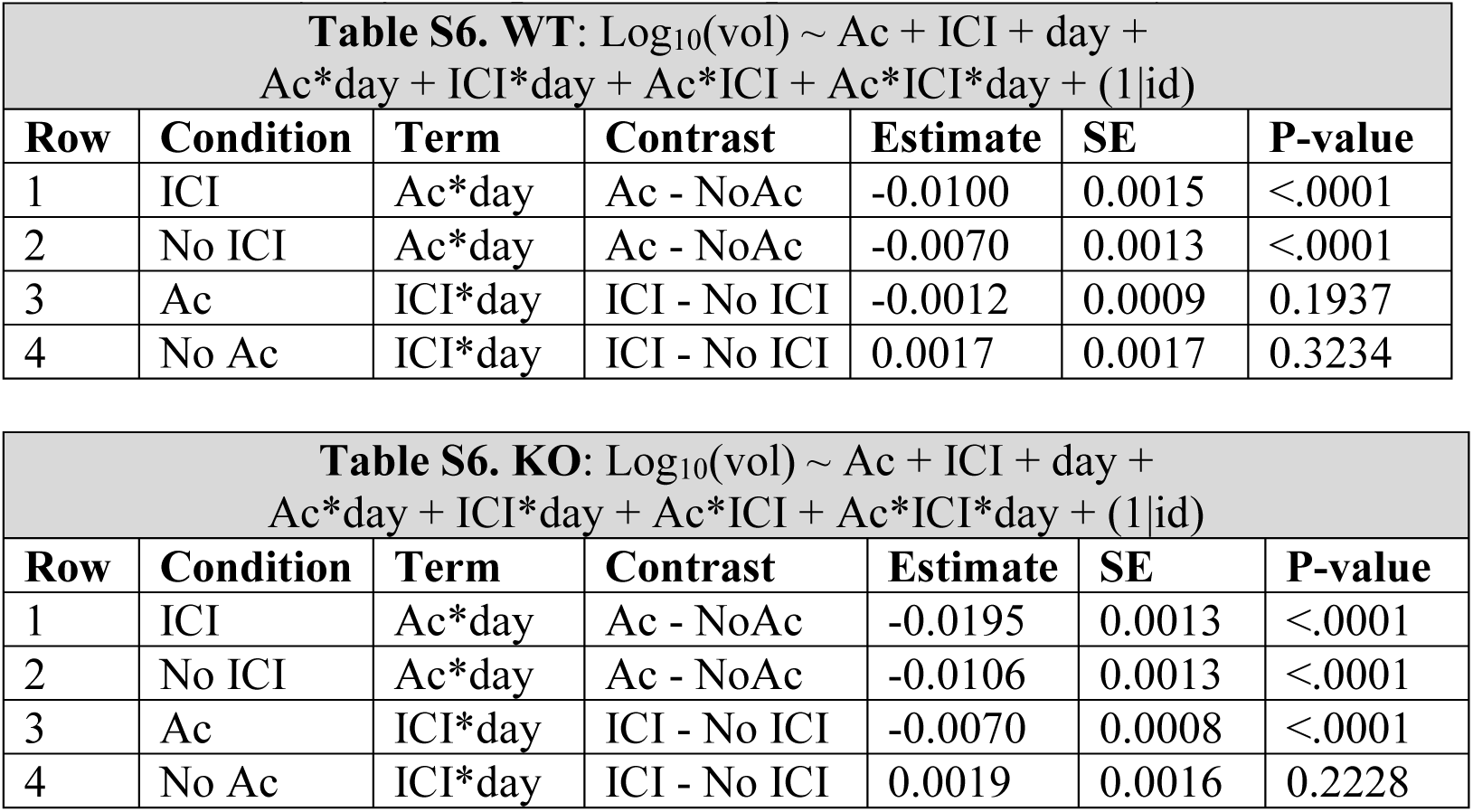
Tukey-adjusted pairwise comparisons of the 3-way interaction

**Table S7.**
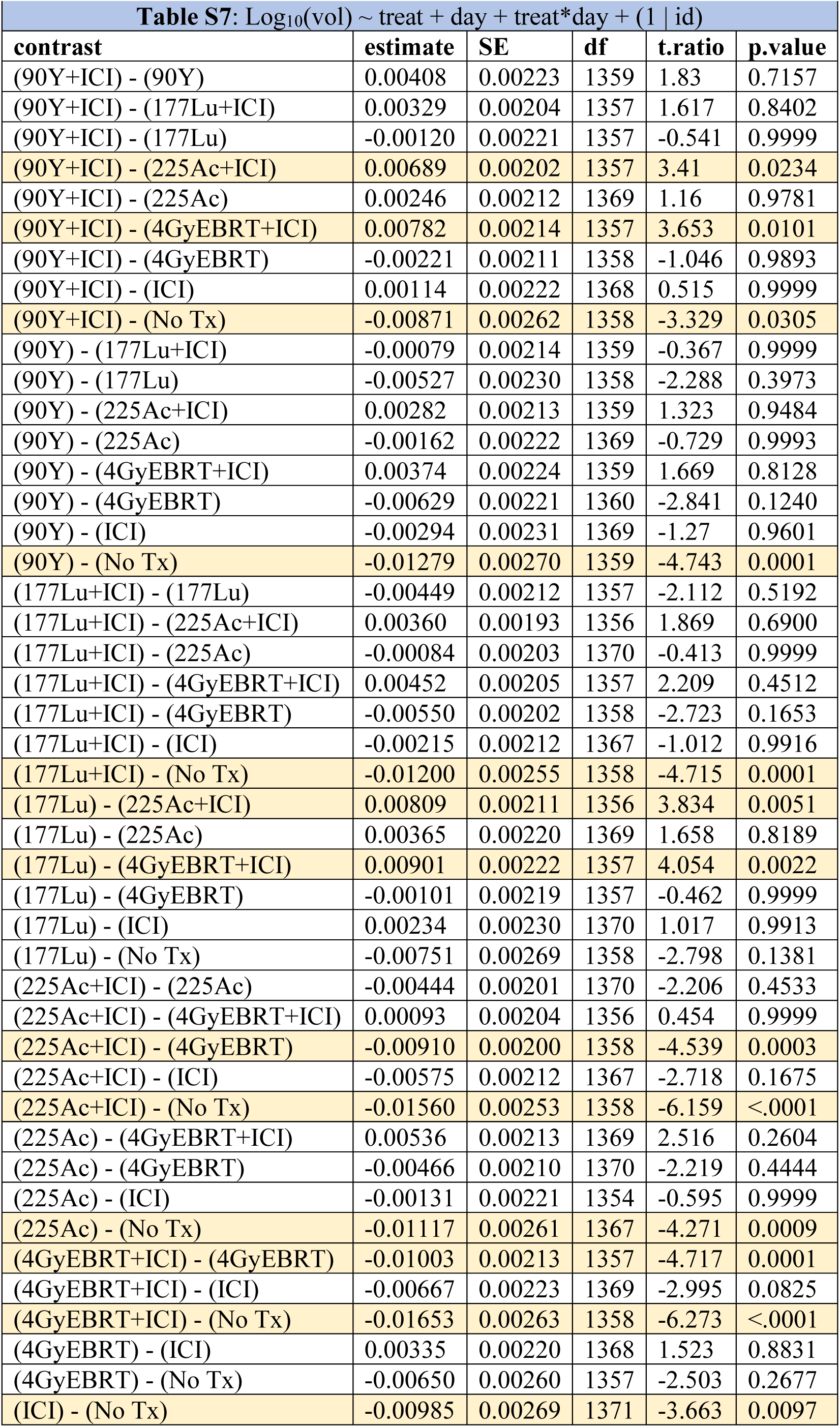
Tukey-adjusted pairwise comparisons of estimated marginal means (tumor growth curves, Figure 7)

**Table S8.**
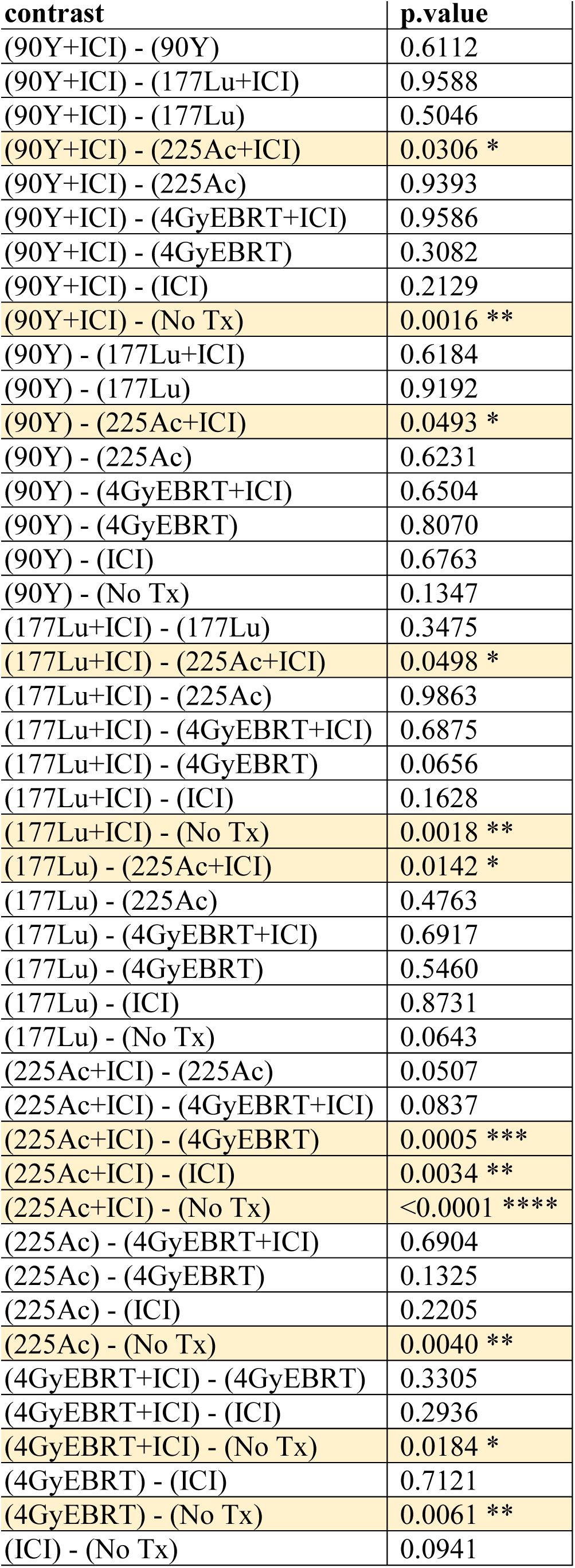
Log-rank (Mantel-Cox) test of overall survival in Figure 7

