## Supplemental Material for "Effects of clinically relevant radionuclides on the activation of a type I interferon response by radiopharmaceuticals in syngeneic murine tumor models"

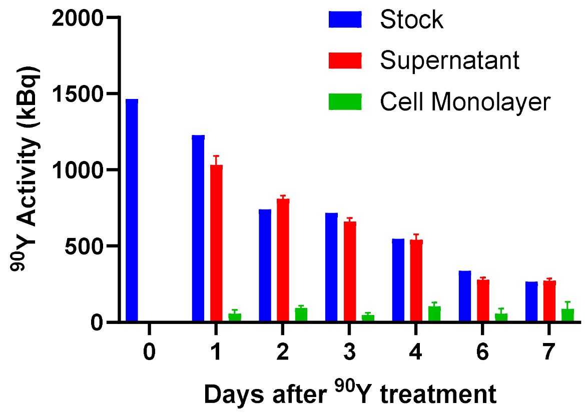


**Fig. S1. Specific activity maintained in radionuclide-containing cell culture media throughout media exchanges.**
1480 kBq/mL $^{90}Y$ in RPMI prepared as a stock solution, and $^{90}Y$ activity in kBq measured with Capintec dose calibrator every 24h between 1-7 days after $^{90}Y$ treatment in 1 mL of: stock solution, cell culture supernatant, or cell monolayer after washing twice with PBS and harvesting by scraping in TRIzol. N=3 for supernatant and cell monolayer, n=1 for stock solution.

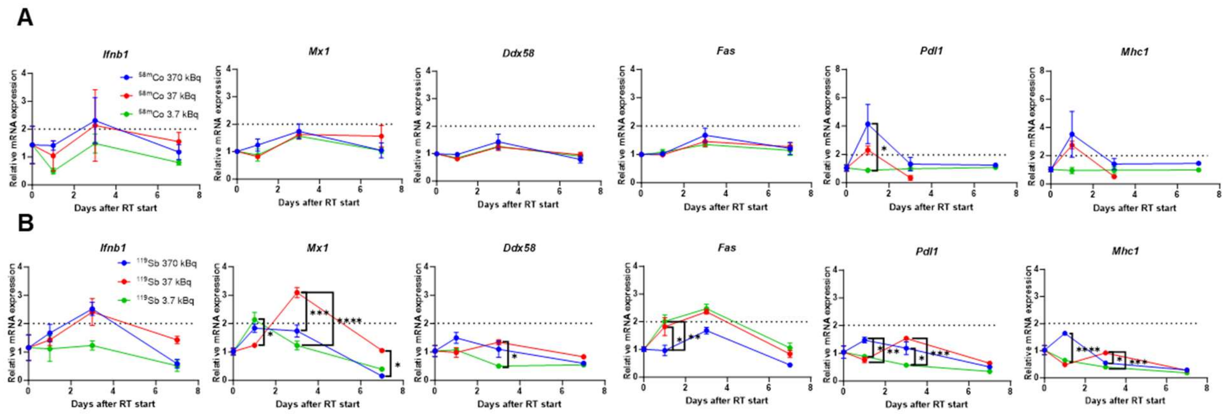


**Fig. S2. Earlier induction of type I IFN response/immunologic response to radiation following $^{58m}$Co and $^{119}$Sb.**

Cells were radiated with either 3.7, 37, or 370 kBq **(A)** $^{58m}$Co or **(B)** $^{119}$Sb and harvested 1, 3, or 7 days following RT. qPCR was used to quantify gene expression and is reported as fold changed normalized to untreated controls. $^{58m}$Co: n=3-6 per treatment group per timepoint; $^{119}$Sb: n=3 per treatment group per timepoint. Two-way ANOVA with Tukey's HSD post hoc test was used to compare fold change in expression between groups.

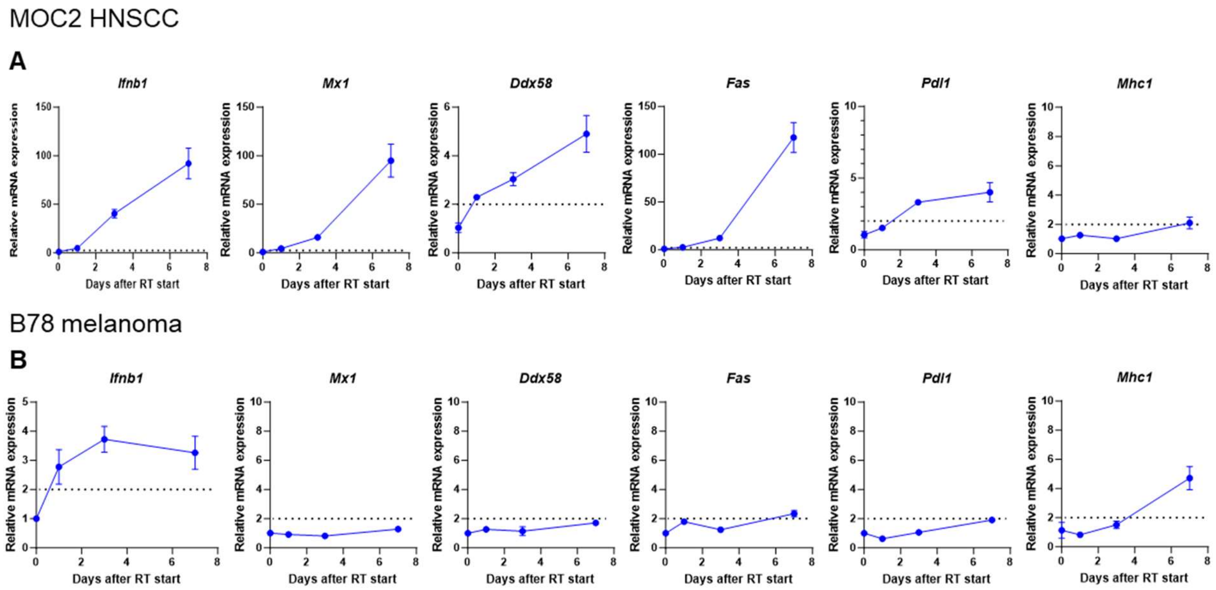


**Fig. S3. Longitudinal *Ifnb1, Mx1, Ddx58, Fas, Pdl1,* and *Mhc1* expression following (A) 12 Gy EBRT (MOC2) or (B) 4 Gy EBRT (B78) *in vitro*.**

Cells were harvested 1, 3, and 7 days following irradiation. qPCR was used to quantify gene expression and is reported as fold changed normalized to untreated controls. MOC2: n=6 per timepoint, B78: n=5-6 per timepoint.

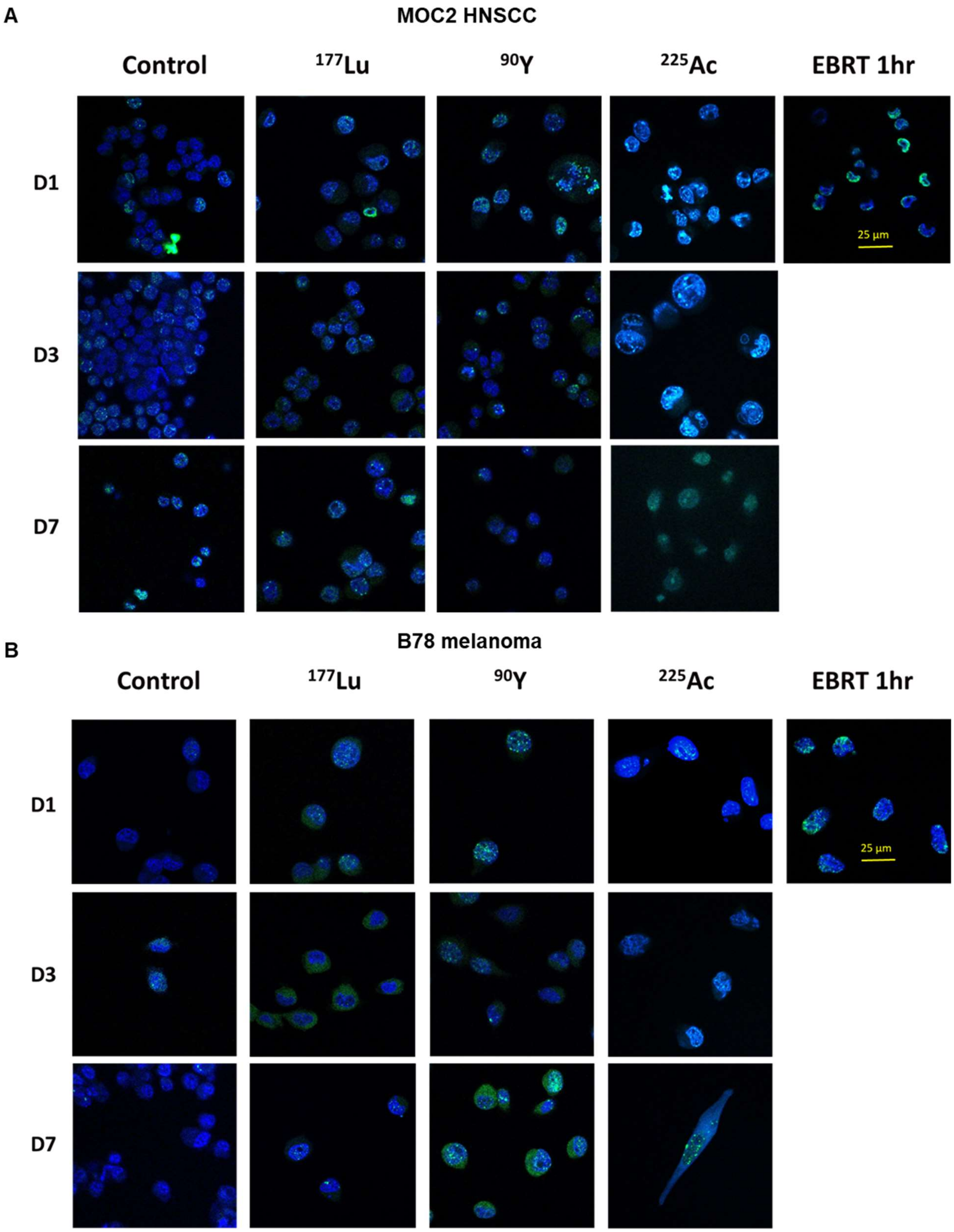


**Fig. S4. Representative images of γH2AX foci over time in (A) MOC2 (12 Gy) and (B) B78 (4 Gy) cells following EBRT, $^{90}$Y, $^{177}$Lu, and $^{225}$Ac.**

Scale bars shown. Images selected from three independent samples per treatment group; 50 cells total per treatment group quantified. γH2AX foci were not quantified for the MOC2 EBRT 1h group due to the high prevalence of apoptotic rings as opposed to nuclear foci nor for the MOC2 $^{225}$Ac day 7 group due to the low number of nuclei present.

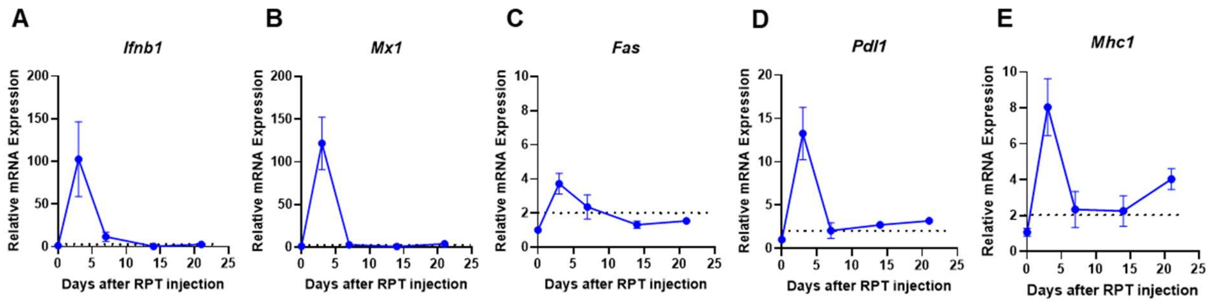


**Fig. S5. Longitudinal *in vivo* gene expression following 12 Gy EBRT (MOC2).**
MOC2 tumors were grown to ~200 mm$^3$ and radiated with 12 Gy tumor targeted EBRT. Tumors were harvested 3, 7, 14, or 21 days following RT administration. **(A-E)** qPCR was used to quantify gene expression and is reported as fold changed normalized to untreated controls. N=3-5 mice per treatment group per timepoint.

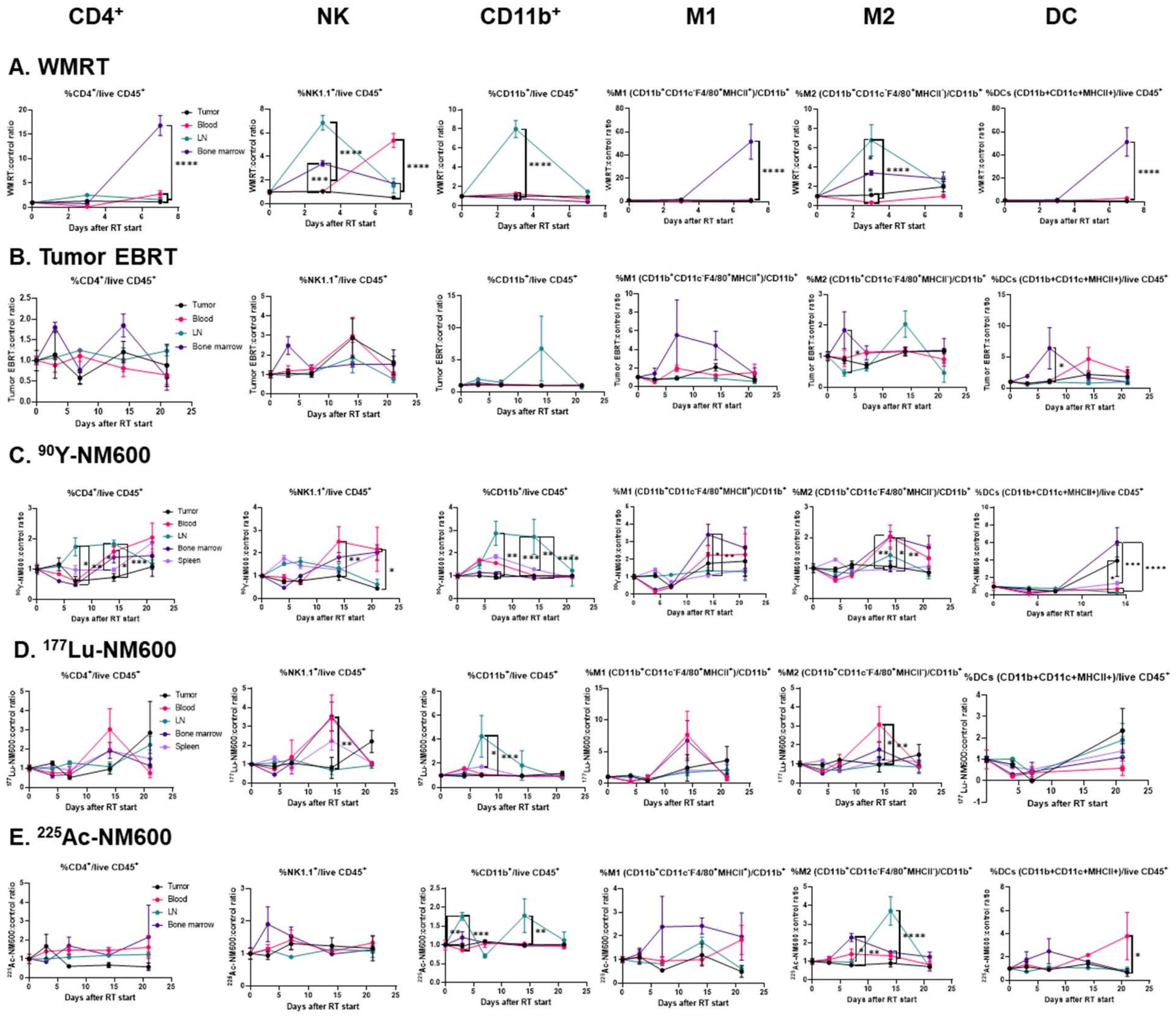


**Fig. S6. $^{90}$Y-, $^{177}$Lu, $^{225}$Ac-NM600 modulate tumor immune cell composition in the tumor microenvironment.**

**(A-E)** From same mice as **Figure 4**, flow cytometry analyses of tumor, blood, tumor-draining lymph node, bone marrow, and spleen immune cell infiltrates [$CD4^+$ T cells, NK cells ($NK1.1^+$), myeloid cells ($CD11b^+$), M1-like macrophages ($CD11b^+CD11c^-F4/80^+MHCII^+$), M2-like macrophages ($CD11b^+CD11c^-F4/80^+MHCII^-$), and dendritic cells ($CD11b^+CD11c^+MHCII^+$)] as a percent of total live cells normalized to mean of control (no treatment) is shown at 3, 7, 14, and 21 days after RT administration in MOC2 HNSCC. N=4-6 per treatment group per timepoint. Two-way ANOVA with Tukey's HSD post hoc test was used to compare ratios between tissue types.

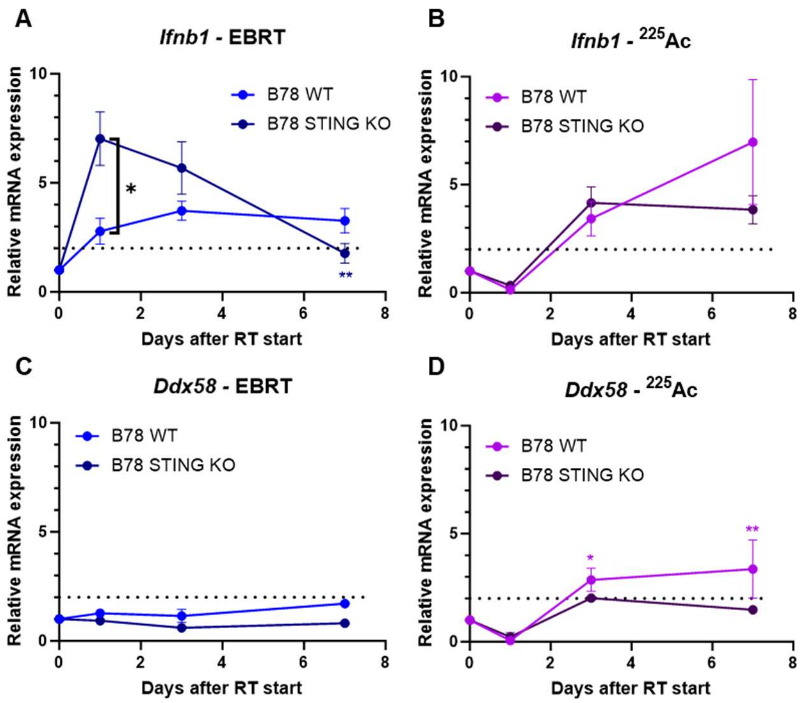


**Fig. S7. B78 WT vs. STING KO cell line in vitro *Ifnb1* and *Ddx58* gene expression upregulation following $^{225}$Ac or EBRT.**
**(A-D)** *In vitro* qPCR following 4 Gy EBRT or $^{225}$Ac. qPCR was used to quantify gene expression and is reported as fold changed normalized to untreated controls. qPCR: n=4-6 per treatment group per timepoint. Two-way ANOVA with Tukey's HSD post hoc test was used to compare fold change in expression between cell lines. Colored asterisks correspond to comparisons to day 1 for EBRT (blue) or $^{225}$Ac (purple). Black asterisks correspond to comparisons between cell lines at the indicated timepoint.

**Table S1. Initial radionuclide activity (kBq) required per milliliter of media as a function of mean absorbed dose (Gy) to cell monolayer at an infinite timepoint**

| Absorbed dose to cell layer (at infinity) | Activity concentration (kBq/mL) | | |
|---|---|---|---|
| | $^{90}$Y | $^{177}$Lu | $^{225}$Ac |
| 4 Gy | 222 | 536.5 | 2.4975 |
| 12 Gy | 647.5 | 1572.5 | 7.4 |

**Table S2. EBRT regimens for *in vitro* studies**

| EBRT regimen | Radionuclide dose at infinity (Gy) | Cell harvest timepoint (days after RT) | RT dose(s) (Gy) | Day(s) RT administered |
|---|---|---|---|---|
| $^{90}$Y EBRT dose equivalent (EBRT-eq) | 12 | 1 | 2.74 | 0 |
| | | 3 | 6.48 | 0 |
| | | 7 | 10.04 | 0 |
| $^{90}$Y EBRT dose fractionation (EBRT-fr) | 12 | 1 | 2.74 | 0 |
| | | 3 | 2.74, 2.11, 1.63 | 0, 1, 2 |
| | | 7 | 2.74, 2.11, 1.63, 1.26, 0.97, 0.75, 0.58 | 0, 1, 2, 3, 4, 5, 6 |
| $^{177}$Lu EBRT-eq | 12 | 1 | 1.19 | 0 |
| | | 3 | 3.22 | 0 |
| | | 7 | 6.23 | 0 |
| $^{177}$Lu EBRT-fr | 12 | 1 | 1.19 | 0 |
| | | 3 | 1.19, 1.07, 0.96 | 0, 1, 2 |
| | | 7 | 1.19, 1.07, 0.96, 0.87, 0.78, 0.71, 0.64 | 0, 1, 2, 3, 4, 5, 6 |
| $^{225}$Ac EBRT-eq | 12 | 1 | 0.81 | 0 |
| | | 3 | 2.27 | 0 |
| | | 7 | 4.64 | 0 |
| $^{225}$Ac EBRT-fr | 12 | 1 | 0.81 | 0 |
| | | 3 | 0.81, 0.76, 0.70 | 0, 1, 2 |
| | | 7 | 0.81, 0.76, 0.70, 0.66, 0.61, 0.57, 0.53 | 0, 1, 2, 3, 4, 5, 6 |
| $^{225}$Ac EBRT-eq | 4 | 1 | 0.27 | 0 |
| | | 3 | 0.76 | 0 |
| | | 7 | 1.55 | 0 |
| $^{225}$Ac EBRT-fr | 4 | 1 | 0.27 | 0 |
| | | 3 | 0.27, 0.25, 0.23 | 0, 1, 2 |
| | | 7 | 0.27, 0.25, 0.23, 0.22, 0.20, 0.19, 0.18 | 0, 1, 2, 3, 4, 5, 6 |

**Table S3. List of TaqMan probes utilized for quantitative RT-PCR experiments**

| Gene Name | Assay ID |
|---|---|
| *Hprt* | Mm03024075_m1 |
| *Ifnb1* | Mm00439552_s1 |
| *Mx1* | Mm00487796_m1 |
| *Fas* | Mm01204974_m1 |
| *Pdl1* (CD274) | Mm03048248_m1 |
| *Mhc1* (H2-D) | Mm04208017_mH |
| *Ddx58* | Mm01216853_m1 |

**Table S4. List of flow cytometry antibody targets, clones, and fluorophores**

| Name | Clone | Fluorophore | Catalog number |
|---|---|---|---|
| CD4 | RM4-5 | FITC | BioLegend 100510 |
| CD11c | N418 | PerCP-Cy5.5 | BioLegend 117328 |
| F4/80 | BM8 | PE/Dazzle 594 | BioLegend 123146 |
| NK1.1 | PK136 | PE-Cy5 | BioLegend 108716 |
| FOXP3 | FJK-16s | PE-Cy7 | Thermo Scientific 25-5773-82 |
| CD279 (PD-1) | RMP1-30 | BV421 | BioLegend 109121 |
| I-A/I-E (MHC-II) | M5/114 | BV605 | BioLegend 107639 |
| CD45 | 30-F11 | BV510 | BioLegend 103138 |
| CD11b | M1/70 | BV711 | BioLegend 101242 |
| CD25 | PC61 | APC | BioLegend 102012 |
| CD8a | 53-6.7 | Alexa700 | BioLegend 100730 |

**Table S5. Tukey-adjusted pairwise comparisons of estimated marginal means**

| **Table S5. Ac+ICI**: $Log_{10}$(vol) ~ line + day + line*day + (1 \| id) | | | | | |
|---|---|---|---|---|---|
| **contrast** | **estimate** | **SE** | **df** | **t.ratio** | **p.value** |
| (STINGKO) - (B78 WT) | 0.0058 | 0.0011 | 267 | 5.424 | <.0001 |

| **Table S5. Ac**: $Log_{10}$(vol) ~ line + day + line*day + (1 \| id) | | | | | |
|---|---|---|---|---|---|
| **contrast** | **estimate** | **SE** | **df** | **t.ratio** | **p.value** |
| (STINGKO) - (B78 WT) | 4.53e-7 | 0.0009 | 248 | 0.001 | 0.9996 |

| **Table S5. ICI**: $Log_{10}$(vol) ~ line + day + line*day + (1 \| id) | | | | | |
|---|---|---|---|---|---|
| **contrast** | **estimate** | **SE** | **df** | **t.ratio** | **p.value** |
| (STINGKO) - (B78 WT) | -0.0038 | 0.0016 | 172 | -2.331 | 0.0209 |

| **Table S5. No Tx**: $Log_{10}$(vol) ~ line + day + line*day + (1 \| id) | | | | | |
|---|---|---|---|---|---|
| **contrast** | **estimate** | **SE** | **df** | **t.ratio** | **p.value** |
| (STINGKO) - (B78 WT) | -0.0037 | 0.0015 | 183 | -2.407 | 0.0171 |

**Table S6. Tukey-adjusted pairwise comparisons of the 3-way interaction**

| **Table S6. WT**: $Log_{10}$(vol) ~ Ac + ICI + day + Ac*day + ICI*day + Ac*ICI + Ac*ICI*day + (1\|id) | | | | | | |
|---|---|---|---|---|---|---|
| **Row** | **Condition** | **Term** | **Contrast** | **Estimate** | **SE** | **P-value** |
| 1 | ICI | Ac*day | Ac - NoAc | -0.0100 | 0.0015 | <.0001 |
| 2 | No ICI | Ac*day | Ac - NoAc | -0.0070 | 0.0013 | <.0001 |
| 3 | Ac | ICI*day | ICI - No ICI | -0.0012 | 0.0009 | 0.1937 |
| 4 | No Ac | ICI*day | ICI - No ICI | 0.0017 | 0.0017 | 0.3234 |

| **Table S6. KO**: $Log_{10}$(vol) ~ Ac + ICI + day + Ac*day + ICI*day + Ac*ICI + Ac*ICI*day + (1\|id) | | | | | | |
|---|---|---|---|---|---|---|
| **Row** | **Condition** | **Term** | **Contrast** | **Estimate** | **SE** | **P-value** |
| 1 | ICI | Ac*day | Ac - NoAc | -0.0195 | 0.0013 | <.0001 |
| 2 | No ICI | Ac*day | Ac - NoAc | -0.0106 | 0.0013 | <.0001 |
| 3 | Ac | ICI*day | ICI - No ICI | -0.0070 | 0.0008 | <.0001 |
| 4 | No Ac | ICI*day | ICI - No ICI | 0.0019 | 0.0016 | 0.2228 |

**Table S7. Tukey-adjusted pairwise comparisons of estimated marginal means (tumor growth curves, Figure 7)**

| **Table S7**: $Log_{10}$(vol) ~ treat + day + treat*day + (1 \| id) | | | | | |
|---|---|---|---|---|---|
| **contrast** | **estimate** | **SE** | **df** | **t.ratio** | **p.value** |
| (90Y+ICI) - (90Y) | 0.00408 | 0.00223 | 1359 | 1.83 | 0.7157 |
| (90Y+ICI) - (177Lu+ICI) | 0.00329 | 0.00204 | 1357 | 1.617 | 0.8402 |
| (90Y+ICI) - (177Lu) | -0.00120 | 0.00221 | 1357 | -0.541 | 0.9999 |
| (90Y+ICI) - (225Ac+ICI) | 0.00689 | 0.00202 | 1357 | 3.41 | 0.0234 |
| (90Y+ICI) - (225Ac) | 0.00246 | 0.00212 | 1369 | 1.16 | 0.9781 |
| (90Y+ICI) - (4GyEBRT+ICI) | 0.00782 | 0.00214 | 1357 | 3.653 | 0.0101 |
| (90Y+ICI) - (4GyEBRT) | -0.00221 | 0.00211 | 1358 | -1.046 | 0.9893 |
| (90Y+ICI) - (ICI) | 0.00114 | 0.00222 | 1368 | 0.515 | 0.9999 |
| (90Y+ICI) - (No Tx) | -0.00871 | 0.00262 | 1358 | -3.329 | 0.0305 |
| (90Y) - (177Lu+ICI) | -0.00079 | 0.00214 | 1359 | -0.367 | 0.9999 |
| (90Y) - (177Lu) | -0.00527 | 0.00230 | 1358 | -2.288 | 0.3973 |
| (90Y) - (225Ac+ICI) | 0.00282 | 0.00213 | 1359 | 1.323 | 0.9484 |
| (90Y) - (225Ac) | -0.00162 | 0.00222 | 1369 | -0.729 | 0.9993 |
| (90Y) - (4GyEBRT+ICI) | 0.00374 | 0.00224 | 1359 | 1.669 | 0.8128 |
| (90Y) - (4GyEBRT) | -0.00629 | 0.00221 | 1360 | -2.841 | 0.1240 |
| (90Y) - (ICI) | -0.00294 | 0.00231 | 1369 | -1.27 | 0.9601 |
| (90Y) - (No Tx) | -0.01279 | 0.00270 | 1359 | -4.743 | 0.0001 |
| (177Lu+ICI) - (177Lu) | -0.00449 | 0.00212 | 1357 | -2.112 | 0.5192 |
| (177Lu+ICI) - (225Ac+ICI) | 0.00360 | 0.00193 | 1356 | 1.869 | 0.6900 |
| (177Lu+ICI) - (225Ac) | -0.00084 | 0.00203 | 1370 | -0.413 | 0.9999 |
| (177Lu+ICI) - (4GyEBRT+ICI) | 0.00452 | 0.00205 | 1357 | 2.209 | 0.4512 |
| (177Lu+ICI) - (4GyEBRT) | -0.00550 | 0.00202 | 1358 | -2.723 | 0.1653 |
| (177Lu+ICI) - (ICI) | -0.00215 | 0.00212 | 1367 | -1.012 | 0.9916 |
| (177Lu+ICI) - (No Tx) | -0.01200 | 0.00255 | 1358 | -4.715 | 0.0001 |
| (177Lu) - (225Ac+ICI) | 0.00809 | 0.00211 | 1356 | 3.834 | 0.0051 |
| (177Lu) - (225Ac) | 0.00365 | 0.00220 | 1369 | 1.658 | 0.8189 |
| (177Lu) - (4GyEBRT+ICI) | 0.00901 | 0.00222 | 1357 | 4.054 | 0.0022 |
| (177Lu) - (4GyEBRT) | -0.00101 | 0.00219 | 1357 | -0.462 | 0.9999 |
| (177Lu) - (ICI) | 0.00234 | 0.00230 | 1370 | 1.017 | 0.9913 |
| (177Lu) - (No Tx) | -0.00751 | 0.00269 | 1358 | -2.798 | 0.1381 |
| (225Ac+ICI) - (225Ac) | -0.00444 | 0.00201 | 1370 | -2.206 | 0.4533 |
| (225Ac+ICI) - (4GyEBRT+ICI) | 0.00093 | 0.00204 | 1356 | 0.454 | 0.9999 |
| (225Ac+ICI) - (4GyEBRT) | -0.00910 | 0.00200 | 1358 | -4.539 | 0.0003 |
| (225Ac+ICI) - (ICI) | -0.00575 | 0.00212 | 1367 | -2.718 | 0.1675 |
| (225Ac+ICI) - (No Tx) | -0.01560 | 0.00253 | 1358 | -6.159 | <.0001 |
| (225Ac) - (4GyEBRT+ICI) | 0.00536 | 0.00213 | 1369 | 2.516 | 0.2604 |
| (225Ac) - (4GyEBRT) | -0.00466 | 0.00210 | 1370 | -2.219 | 0.4444 |
| (225Ac) - (ICI) | -0.00131 | 0.00221 | 1354 | -0.595 | 0.9999 |
| (225Ac) - (No Tx) | -0.01117 | 0.00261 | 1367 | -4.271 | 0.0009 |
| (4GyEBRT+ICI) - (4GyEBRT) | -0.01003 | 0.00213 | 1357 | -4.717 | 0.0001 |
| (4GyEBRT+ICI) - (ICI) | -0.00667 | 0.00223 | 1369 | -2.995 | 0.0825 |
| (4GyEBRT+ICI) - (No Tx) | -0.01653 | 0.00263 | 1358 | -6.273 | <.0001 |
| (4GyEBRT) - (ICI) | 0.00335 | 0.00220 | 1368 | 1.523 | 0.8831 |
| (4GyEBRT) - (No Tx) | -0.00650 | 0.00260 | 1357 | -2.503 | 0.2677 |
| (ICI) - (No Tx) | -0.00985 | 0.00269 | 1371 | -3.663 | 0.0097 |

**Table S8. Log-rank (Mantel-Cox) test of overall survival in Figure 7**

| **Table S8**: Kaplan-Meier Survival Log-rank test | |
|---|---|
| **contrast** | **p.value** |
| (90Y+ICI) - (90Y) | 0.6112 |
| (90Y+ICI) - (177Lu+ICI) | 0.9588 |
| (90Y+ICI) - (177Lu) | 0.5046 |
| (90Y+ICI) - (225Ac+ICI) | 0.0306 * |
| (90Y+ICI) - (225Ac) | 0.9393 |
| (90Y+ICI) - (4GyEBRT+ICI) | 0.9586 |
| (90Y+ICI) - (4GyEBRT) | 0.3082 |
| (90Y+ICI) - (ICI) | 0.2129 |
| (90Y+ICI) - (No Tx) | 0.0016 ** |
| (90Y) - (177Lu+ICI) | 0.6184 |
| (90Y) - (177Lu) | 0.9192 |
| (90Y) - (225Ac+ICI) | 0.0493 * |
| (90Y) - (225Ac) | 0.6231 |
| (90Y) - (4GyEBRT+ICI) | 0.6504 |
| (90Y) - (4GyEBRT) | 0.8070 |
| (90Y) - (ICI) | 0.6763 |
| (90Y) - (No Tx) | 0.1347 |
| (177Lu+ICI) - (177Lu) | 0.3475 |
| (177Lu+ICI) - (225Ac+ICI) | 0.0498 * |
| (177Lu+ICI) - (225Ac) | 0.9863 |
| (177Lu+ICI) - (4GyEBRT+ICI) | 0.6875 |
| (177Lu+ICI) - (4GyEBRT) | 0.0656 |
| (177Lu+ICI) - (ICI) | 0.1628 |
| (177Lu+ICI) - (No Tx) | 0.0018 ** |
| (177Lu) - (225Ac+ICI) | 0.0142 * |
| (177Lu) - (225Ac) | 0.4763 |
| (177Lu) - (4GyEBRT+ICI) | 0.6917 |
| (177Lu) - (4GyEBRT) | 0.5460 |
| (177Lu) - (ICI) | 0.8731 |
| (177Lu) - (No Tx) | 0.0643 |
| (225Ac+ICI) - (225Ac) | 0.0507 |
| (225Ac+ICI) - (4GyEBRT+ICI) | 0.0837 |
| (225Ac+ICI) - (4GyEBRT) | 0.0005 *** |
| (225Ac+ICI) - (ICI) | 0.0034 ** |
| (225Ac+ICI) - (No Tx) | <0.0001 **** |
| (225Ac) - (4GyEBRT+ICI) | 0.6904 |
| (225Ac) - (4GyEBRT) | 0.1325 |
| (225Ac) - (ICI) | 0.2205 |
| (225Ac) - (No Tx) | 0.0040 ** |
| (4GyEBRT+ICI) - (4GyEBRT) | 0.3305 |
| (4GyEBRT+ICI) - (ICI) | 0.2936 |
| (4GyEBRT+ICI) - (No Tx) | 0.0184 * |
| (4GyEBRT) - (ICI) | 0.7121 |
| (4GyEBRT) - (No Tx) | 0.0061 ** |
| (ICI) - (No Tx) | 0.0941 |
